# A metabolic checkpoint coordinates bacterial cell envelope biosynthesis and c-di-GMP signaling

**DOI:** 10.64898/2026.06.11.731730

**Authors:** Xuhui Zheng, Katayoun Daneshjoo, Angeli Shieh, Megan O’Malley, Aashi Shah, Matthew Parsek

**Affiliations:** Department of Microbiology, University of Washington, Seattle, WA, USA

## Abstract

The bacterial second messenger cyclic dimeric guanosine monophosphate (c-di-GMP) drives the transition from motile to biofilm lifestyles, yet the mechanisms by which bacteria couple cell envelope stress to c-di-GMP signaling remain poorly understood. Here, we show that sub-inhibitory concentrations of antibiotics targeting early cytoplasmic steps of peptidoglycan (PG) biosynthesis, but not inhibitors of PG polymerization, membrane integrity, or other intracellular processes, specifically elevate intracellular c-di-GMP levels in *Pseudomonas aeruginosa*. Using live-cell imaging and *in vitro* enzymatic assays, we demonstrate that this elevation results from reduced phosphodiesterase (PDE) activity rather than increased diguanylate cyclase (DGC) activity, with multiple PDEs contributing to the response. A screen of the complete *P. aeruginosa* DGC/PDE mutant library identified DipA, BifA, RocR, and RmcA as the primary PDEs mediating this effect. Strikingly, we find that acetyl-CoA, a central metabolite consumed during cell envelope precursor biosynthesis, directly inhibits PDE activity by competing for the conserved EAL domain active site, as supported by biochemical assays with purified RocR and molecular docking analysis. Because EAL domain residues that contact acetyl-CoA are broadly conserved across bacterial species, this mechanism may represent a widespread strategy for sensing metabolic perturbations of cell envelope synthesis. Together, these findings reveal that *P. aeruginosa* monitors the metabolic status of cell envelope biogenesis through acetyl-CoA-mediated allosteric inhibition of c-di-GMP PDEs, linking envelope biosynthetic flux to adaptive biofilm formation.

## INTRODUCTION

Signaling through cyclic diguanylate monophosphate (c-di-GMP) is critical for biofilm formation in a wide range of bacterial species. Upon c-di-GMP production, cells reduce flagellar motility and synthesize an extracellular matrix, promoting the development of cellular aggregates(1). This aggregated lifestyle protects bacteria from multiple environmental stressors, including harsh environmental conditions, host immune responses, and antimicrobial treatments(2, 3). C-di-GMP is synthesized from GTP by diguanylate cyclases (DGCs), and degraded into GMP, either directly or via the intermediate pGpG, by phosphodiesterases (PDEs). In the opportunistic pathogen *Pseudomonas aeruginosa*, a model organism for biofilm studies, c-di-GMP levels are regulated by more than 40 DGCs or PDEs(4). Many of these proteins contain both GGDEF and EAL domains, suggesting the potential for dual DGC/PDE functionality. The complex regulation of c-di-GMP highlights its importance, enabling bacteria to respond to diverse environmental stimuli and transition to a protective growth mode when confronted with a potentially hostile environment.

The cell envelope serves as the first line of defense for bacteria under environmental stress. In Gram-negative bacteria, the envelope typically consists of an inner phospholipid membrane, a thin peptidoglycan (PG) layer, and an outer membrane composed of lipopolysaccharides (LPS) in the outer leaflet and phospholipids in the inner leaflet. The cell envelope is essential for bacterial survival, serving to maintain cell shape, withstand turgor pressure, enable nutrient transport, and support surface colonization(5–7). Owing to its unique structure and critical functions, the bacterial cell envelope has long been a major target for antimicrobial therapies(8).

To maintain cell envelope homeostasis, bacteria employ multiple regulatory systems that respond to cell envelope stress (reviewed in (9, 10)). These systems typically modulate the biosynthesis of envelope components, facilitating repair and quality control. It has also become evident that cell envelope stress elicits broader effects on the cell, including the promotion of biofilm formation(11–13). In addition, membrane-fluidizing agents such as SDS, ethanol, and glycerol have been shown to increase c-di-GMP levels in *P. aeruginosa*, leading to increased biofilm formation(14, 15). DGCs such as SiaD, WspR, and GcbA have been proposed to mediate this process(15–17). Notably, WspR can also be activated by a surface contact-associated stress signal(18). However, both small organic molecules and surface contact may compromise the cell envelope in multiple ways. The precise nature of these stress signals and the mechanisms by which they stimulate c-di-GMP signaling remain unclear. In particular, it is not known whether cell envelope biosynthetic and turnover intermediates resulting from impaired biogenesis or disrupted envelope components can serve as a signal.

Here, we set out to identify the molecular mechanisms by which cell envelope perturbations are sensed and relayed to the c-di-GMP signaling network. By screening a diverse panel of antimicrobial molecules targeting distinct envelope components at sub-MIC concentrations, we discovered that antibiotics disrupting early cytoplasmic steps of PG biosynthesis specifically elevate intracellular c-di-GMP levels, a selectivity that points to a dedicated sensing mechanism rather than a generic stress response. Using live-cell imaging and *in vitro* enzymatic assays, we demonstrate that this elevation is driven by reduced c-di-GMP PDE activity rather than increased DGC activity, with multiple PDEs contributing to the response. Strikingly, we identify acetyl-CoA, a central metabolite consumed during cell envelope precursor biosynthesis, as a direct inhibitor of PDE activity, likely acting by competing for the conserved EAL domain active site. Because the EAL domain residues predicted to contact acetyl-CoA are broadly conserved across bacterial species, this mechanism may represent a widespread strategy by which bacteria monitor the metabolic status of cell envelope biogenesis and mount a c-di-GMP-dependent adaptive response. These findings reveal an unexpected link between central carbon metabolism and second messenger signaling, with important implications for understanding how bacteria initiate biofilm formation in response to cell envelope-targeting antibiotics.

## RESULTS

### Screening a library of antimicrobial molecules for c-di-GMP activation

To probe how cell envelope stress activates c-di-GMP signaling, we assembled a panel of molecules and antibiotics with defined effects on distinct envelope components (Table S1). These included broad-spectrum envelope disruptors (SDS, ethanol, triton X-100, EDTA, glycerol, urea, H_2_O_2_), agents perturbing membrane function (nigericin, oligomycin, valinomycin, NaCl, benzyl alcohol), inhibitors of peptidoglycan synthesis (fosfomycin, D-cycloserine, tunicamycin, ramoplanin, vancomycin, moenomycin, carbenicillin, ceftazidime, meropenem), LPS-targeting antibiotics (colistin, polymyxin B), inhibitors of fatty acid and phospholipid synthesis (bacitracin, fosmidomycin, triclosan), and the cytoskeleton-targeting drug A22. Representative antibiotics targeting intracellular processes (trimethoprim, ciprofloxacin, kanamycin, tetracycline) were also included as controls. C-di-GMP levels were monitored using a dual-fluorescent reporter plasmid consisting of a mGreenLantern (mGL) driven by the *cdrA* promoter and a constitutive mScarlet-I control under the *rpoD* promoter (Fig. S1). The ASV degradation tag fused to mGL and mScarlet-I reduced their half-lives, allowing dynamic monitoring of reporter activity. To optimize growth conditions for the screen, we compared growth and c-di-GMP reporter activity in wild-type (WT) and a *ΔwspRΔsadCΔsiaD* triple DGC mutant across several defined, low-fluorescent media for *P. aeruginosa*, including FAB supplemented with glucose or glutamate, VBMM, Jensen’s, M9 supplemented with glucose, and PM supplemented with casamino acid (PM+CAA) (Fig. S2-S3). In all media, WT exhibited significantly higher reporter activity than the *ΔwspRΔsadCΔsiaD* mutant, indicating that DGCs are actively producing c-di-GMP in these conditions (Fig. S3). PM+CAA supported the fastest growth (Fig. S2) and was therefore selected for all subsequent experiments.

For each molecule, we performed 2-fold serial dilutions to determine MIC values from both OD600 measurements and constitutively expressed mScarlet-I fluorescence (Fig. 1A; full OD600 curves in Fig. S4). C-di-GMP reporter activity was quantified by normalizing mGL by mScarlet-I fluorescence throughout the growth curve (Fig. S5). Figure 1A summarizes reporter activity, normalized to wild-type, during mid-exponential to early stationary phase. As expected, SDS and ethanol strongly activated c-di-GMP signaling at sub-MIC (Fig. 1A-B). We also found that Triton X-100 and EDTA positively affected c-di-GMP signaling (Fig. 1A-B). Among molecules with defined targets in the cell envelope, only a subset of cell wall-targeting antibiotics increased reporter activity (Fig. 1A): fosfomycin and ramoplanin presented significant and consistent activation, whereas D-cycloserine and tunicamycin showed a positive trend (Fig. 1C). In contrast, carbenicillin, which inhibits peptidoglycan polymerization, and colistin, which targets LPS, had no effect (Fig. 1C). LC-MS analysis confirmed that fosfomycin elevated intracellular c-di-GMP levels, while carbenicillin did not (Fig. 1D).

**Fig. 1.**
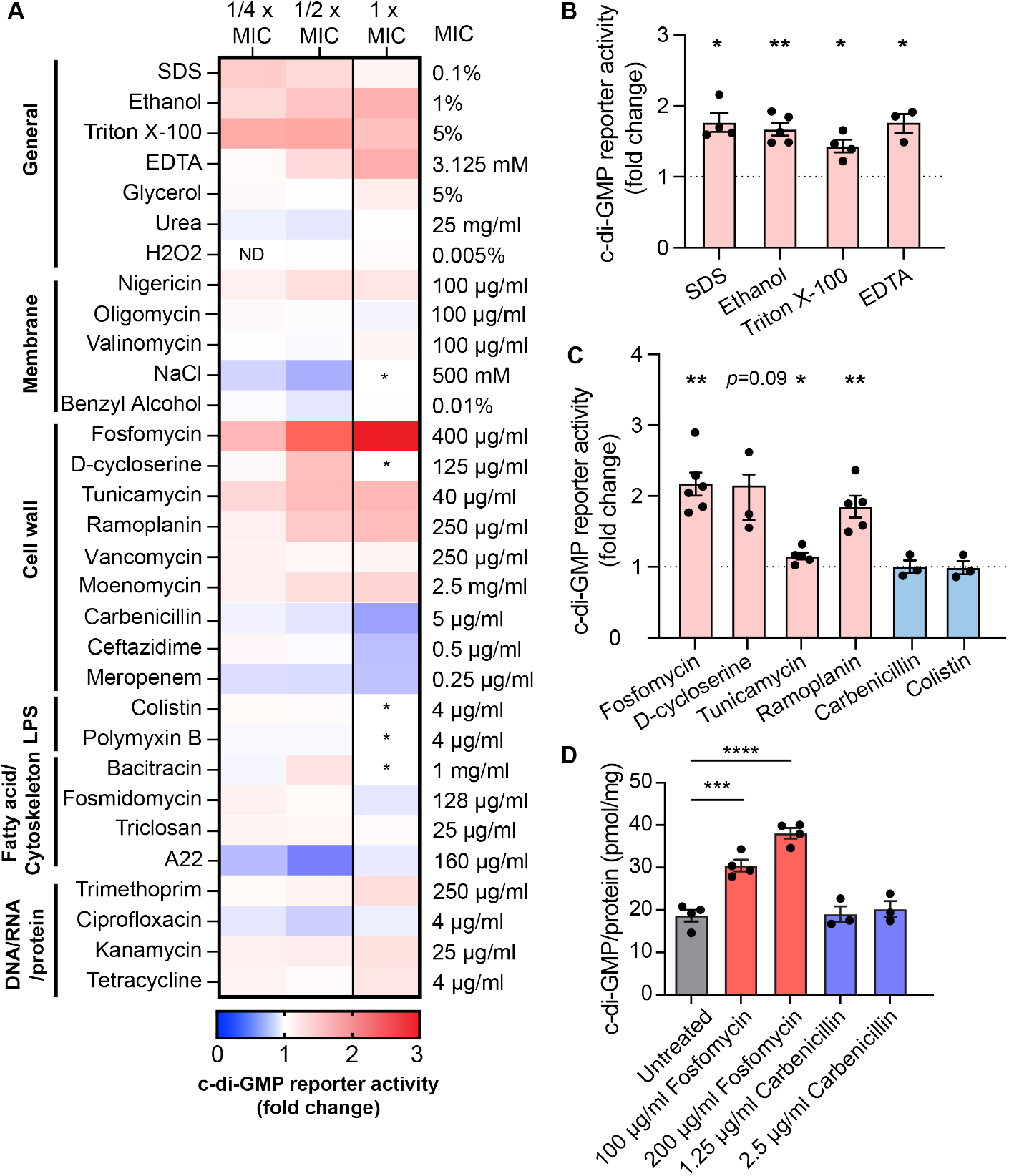
Stressor screen for activation of c-di-GMP signaling. (A) Heatmap of c-di-GMP reporter activity in at 1x, 1/2x, 1/4x MIC of each molecule. Data represent the area under the curve of the c-di-GMP reporter activity from 5 h to 12.5 h of growth, corresponding to mid-exponential to early stationary phase of growth. Values were normalized to the untreated condition within each experiment. ND, not determined. *, the growth defect was too severe to confidently assess reporter activity. (B-C) c-di-GMP reporter activity in the presence of the indicated stressors at 6 h in the growth curve. Values were normalized to the untreated condition within each experiment. Data show mean ± SEM of independent experiments. *, *p < 0*.*05; **, p* < 0.01, determined by one-sample t test against 1 (untreated condition). Stressors concentrations: SDS, 0.0125%; Ethanol, 0.5%; Triton X-100, 2.5%; EDTA, 1.5625 mM; Fosfomycin, 100 µg/ml; D-cycloserine, 62.5 µg/ml; Tunicamycin, 10 µg/ml; Ramoplanin, 125 µg/ml; Carbenicillin, 2.5 µg/ml; Colistin, 2 µg/ml. (D) Quantification of c-di-GMP by LC-MS after growth in the indicated conditions. ***, *p <* 0.001; ****, *p* < 0.0001, determined by ordinary one-way ANOVA with Dunnett’s multiple comparisons test.

Intriguingly, all antibiotics that activated c-di-GMP signaling targeted cytoplasmic steps in PG synthesis: fosfomycin inhibits MurA, D-cycloserine blocks D-alanine incorporation into the pentapeptide, tunicamycin inhibits MraY, and ramoplanin binds to lipid I and lipid II, sequestering their conversions (Fig. 2A, Table S1). In contrast, β-lactam antibiotics targeting the final polymerization steps of PG synthesis had no effect. This pattern suggests that c-di-GMP activation is not a generic consequence of cell envelope stress, but instead reflects sensitivity to disruption of the early cytoplasmic stages of PG biosynthesis. We propose that *P. aeruginosa* coordinates cell envelope synthesis with c-di-GMP production, actively monitoring PG biosynthetic flux and, in response to defects, modulating DGC or PDE activity to elevate intracellular c-di-GMP.

**Fig. 2.**
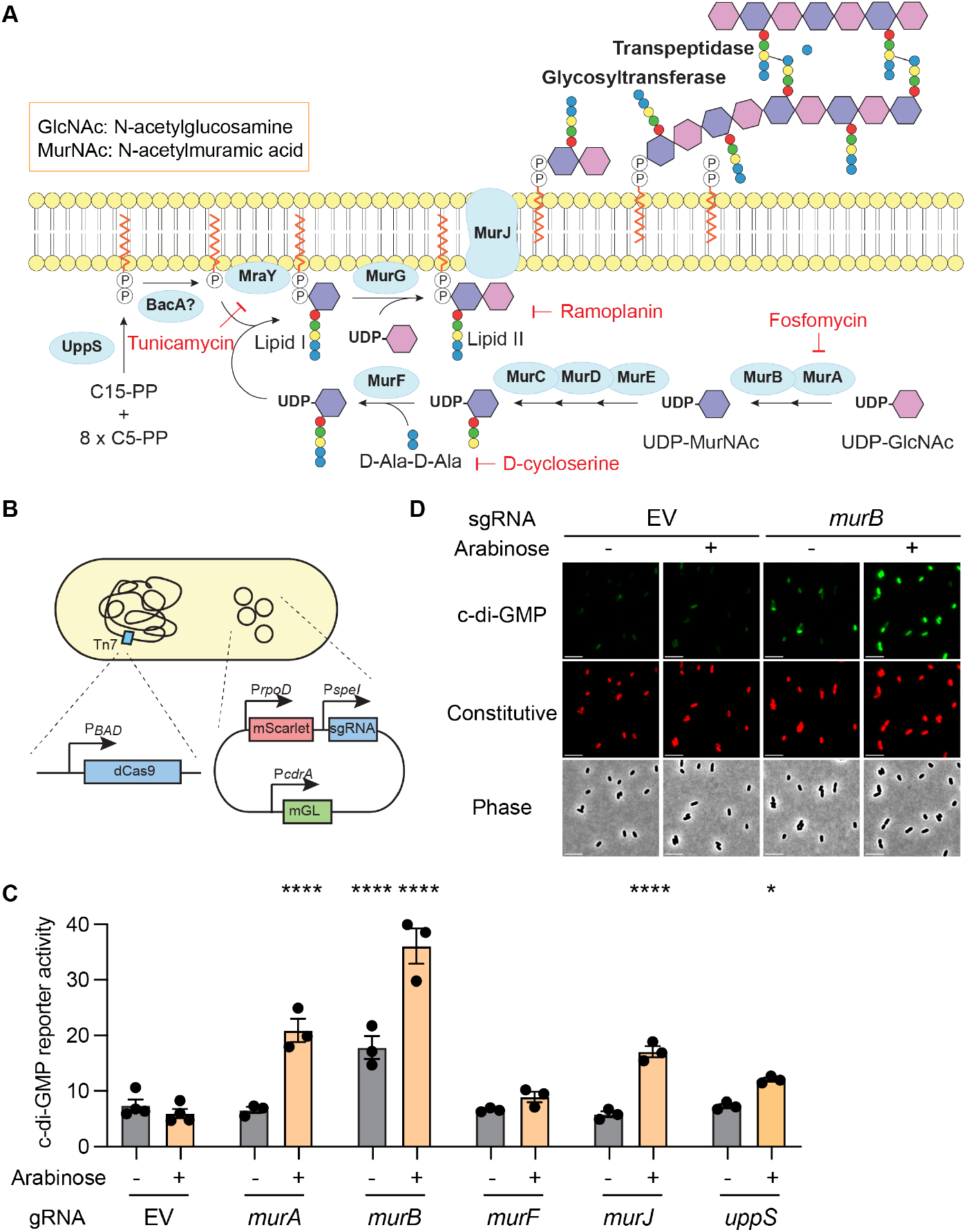
CRISPRi-mediated knockdown of cell wall biosynthesis genes. (A) Schematic of the peptidoglycan synthesis pathway. Enzymes catalyzing each reaction are shown in blue circles. Multiple glycosyltransferases and transpeptidases contribute to peptidoglycan polymerization and crosslinking. (B) Schematic of the CRISPRi design. Deactivated Cas9 (dCas9) is induced by arabinose. The constitutively expressed sgRNA is encoded on a plasmid containing a c-di-GMP-responsive promoter (P*cdrA*) and a constitutive promoter control (P*rpoD*). (C) c-di-GMP reporter activity in CRISPRi strains with sgRNA targeting the indicated genes after 6 h of growth in the presence or absence of arabinose. Arabinose induction of dCas9 enhances CRISPRi knockdown efficiency. Data represent mean ± SEM of independent experiments. \**p* < 0.05, *\*\*\*\*p* < 0.0001 compared with empty vector (EV) in each arabinose condition, determined by two-way ANOVA with Dunnett’s multiple comparisons test. (D) Representative micrographs of murB compared to empty vector control. Scale bar, 5 µm.

### CRISPRi approach to investigate peptidoglycan synthesis

To pinpoint which steps of PG synthesis are monitored by the c-di-GMP machinery, we used a CRISPR interference (CRISPRi) strategy to selectively knock down essential PG biosynthetic enzymes. The reporter plasmid was modified to carry a constitutively expressed single guide RNA (sgRNA) targeting the gene of interest, while a deactivated Cas9 (dCas9) was expressed from an arabinose-inducible promoter at the mini-Tn7 site in the chromosome (Fig. 2B). Upon arabinose induction, dCas9 is recruited to the sgRNA-target complex, blocking transcription. Genes encoding MurF, MurC, MurD, MurE, MraY, and MurG reside in the same operon; thus, an sgRNA against *murF* was expected to inhibit transcription of the entire operon via transcriptional polarity. Additional sgRNAs were designed to target *murA, murB, murJ*, and *uppS*. All sgRNAs were validated by growth inhibition upon arabinose induction (Fig. S6A). Knockdown of *murA, murB*, and *murJ* significantly increased c-di-GMP reporter activity (Fig. 2C). Notably, the *murB* sgRNA triggered elevated reporter activity even without arabinose, suggesting high knockdown efficiency of this sgRNA. Knockdown of *uppS* produced a modest increase, whereas silencing the *murF* operon had no detectable effect (Fig. 2C).

Because growth defects can confound bulk culture measurements of c-di-GMP reporter activity, we also examined reporter activity at the single-cell level by fluorescence microscopy. After equal duration of growth with or without arabinose, the constitutive reporter exhibited uniform fluorescence, indicating comparable levels of protein expression and plasmid copy. In contrast, induction of the *murB* sgRNA markedly increased the proportion of cells with bright c-di-GMP reporter signal (Fig. 2D). This activation was heterogeneous, with a distinct subpopulation retaining low reporter activity despite arabinose induction, suggesting variability in the cellular response to PG synthesis disruption (Fig. S6B).

The contrasting phenotypes resulting from *murF* versus *murA*/*murB* knockdown suggest the existence of a specific checkpoint during cytoplasmic PG synthesis. Notably, the enzymatic steps catalyzed by MurA, MurB, and MurJ are each intimately connected to UDP-GlcNAc incorporation into cell wall precursors. This raises the possibility that fluctuations in UDP-GlcNAc levels act as a metabolic cue, coordinating cell envelope integrity with intracellular signaling pathways. This led us to hypothesize that *P. aeruginosa* monitors UDP-GlcNAc utilization as a sentinel signal, triggering c-di-GMP-mediated adaptations to maintain envelope homeostasis and promote survival under cell wall stress.

### Epistasis analysis of UDP-GlcNAc synthesis and utilization

UDP-GlcNAc is a central metabolic intermediate that links primary carbon metabolism to the synthesis of diverse cell-envelope structures (Fig. 3A). In bacteria, UDP-GlcNAc is synthesized from fructose-6-phosphate through the sequential action of GlmS, GlmM, and GlmU, with GlmU functioning as a bifunctional enzyme possessing both acetyltransferase and uridyltransferase activities(19). Beyond PG biosynthesis, UDP-GlcNAc is also diverted into LPS biosynthesis. The essential enzyme LpxA initiates lipid A synthesis by acylating UDP-GlcNAc. O-antigen assembly begins with transfer of the GlcNAc moiety to the undecaprenyl lipid carrier, a step catalyzed by a variety of nonessential enzymes that account for O-antigen diversity(20). UDP-GlcNAc can also be converted into its epimer UDP-GalNAc, the subunit for the c-di-GMP-regulated exopolysaccharide Pel(21). Since no dedicated UDP-GlcNAc epimerase has been characterized in *P. aeruginosa*, UDP-GalNAc production is thought to occur via the promiscuous activity of UDP-sugar epimerases(22). Together, these pathways position UDP-GlcNAc as a central metabolic hub whose flux is likely monitored by bacteria to modulate c-di-GMP-dependent cell-envelope responses.

**Fig. 3.**
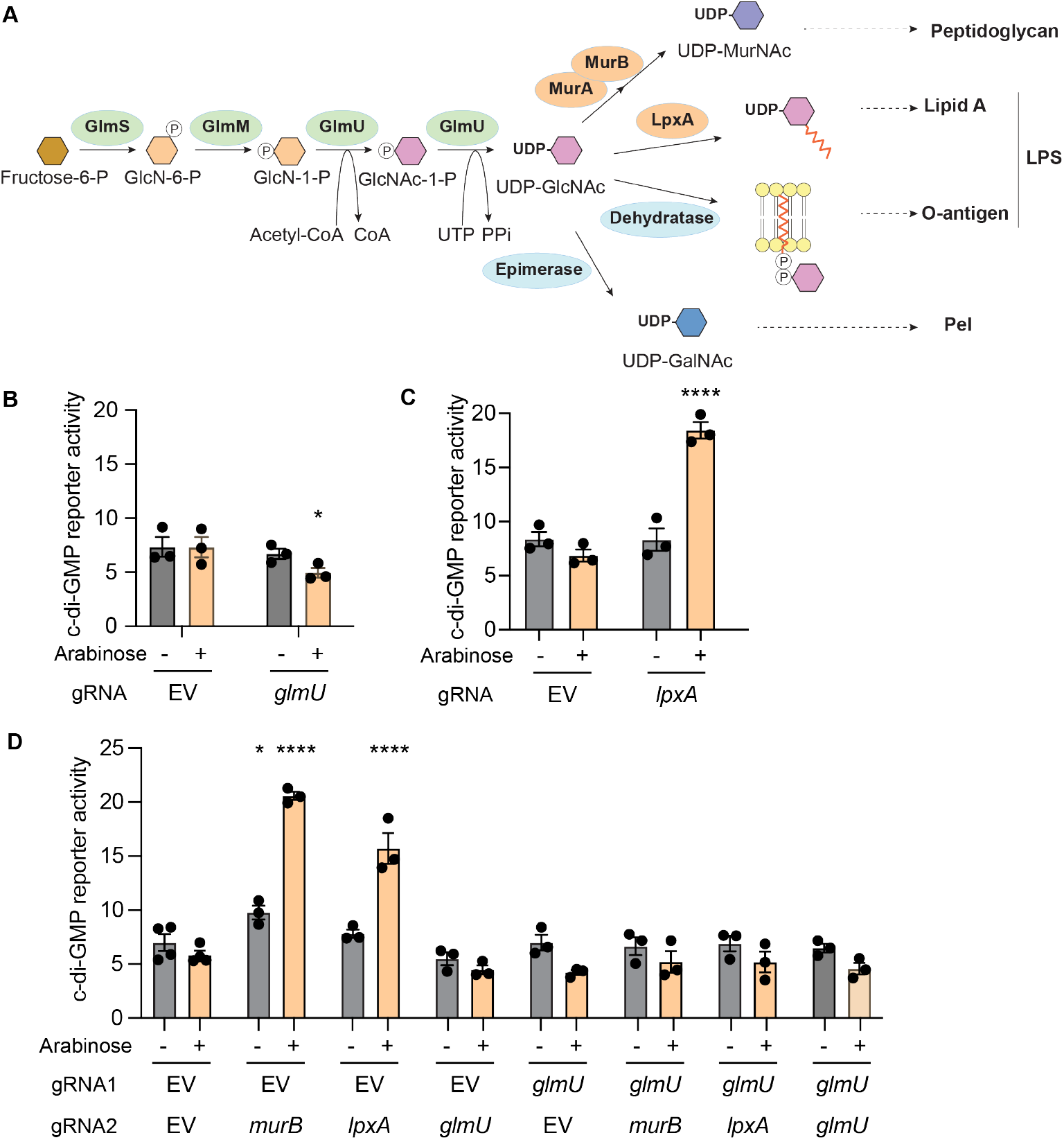
Double CRISPRi-mediated knockdown targeting UDP-GlcNAc metabolism. (A) Schematic of UDP-GlcNAc biosynthesis and its conversion to peptidoglycan, lipid A, O-antigen, and Pel synthesis in *P. aeruginosa*. Enzymes catalyzing each reaction are shown in blue circles. Multiple dehydratases can commit UDP-GlcNAc to O-antigen synthesis. The identity of the epimerase converting UDP-GlcNAc into UDP-GalNAc remains unclear. (B-C) c-di-GMP reporter activity in CRISPRi strains with sgRNA targeting *glmU* or *lpxA* after 6 h of growth in the presence or absence of arabinose to induce dCas9 expression. Data represent mean ± SEM of independent experiments. *\*p* < 0.05, *\*\*p* < 0.01, *\*\*\*\*p* < 0.0001 compared with EV in each arabinose condition, determined by two-way ANOVA with Dunnett’s multiple comparisons test. (D) c-di-GMP reporter activity in strains with two sgRNAs targeting EV or *glmU* in combination with EV, *murB, lpxA*, or *glmU*, after 6 h of growth in the presence or absence of. Data represent mean ± SEM of independent experiments. \**p* < 0.05, *\*\*\*\*p* < 0.0001 compared with EV in each arabinose condition, determined by two-way ANOVA with Dunnett’s multiple comparisons test.

To test the hypothesis that UDP-GlcNAc availability, rather than PG synthesis, serves as the metabolic cue regulating c-di-GMP signaling, we perturbed additional steps in UDP-GlcNAc biosynthesis and utilization. Individual knockdowns of *glmS, glmM*, and *glmU* each reduced c-di-GMP reporter activity (Fig. 3B), indicating that UDP-GlcNAc synthesis is important for c-di-GMP signaling. Notably, although *glmS* and *glmM* knockdowns did not produce a significant growth defect, possibly reflecting lower sgRNA efficiency, a significant reduction in c-di-GMP reporter activity was nonetheless observed. In contrast, knockdown of *lpxA*, which diverts UDP-GlcNAc into lipid A-core biosynthesis, significantly increased c-di-GMP reporter activity (Fig. 3C). Together with the *murA/B* knockdown results, these findings show that blocking UDP-GlcNAc consumption across distinct envelope biogenesis pathways consistently elevates c-di-GMP levels.

To determine whether the *murA/B* and *lpxA* effects are mediated by UDP-GlcNAc, we constructed reporter plasmids expressing two sgRNAs simultaneously. While single knockdowns of *murB* or *lpxA* increased c-di-GMP reporter activity, co-expression with *glmU* sgRNA abolished this increase, yielding activity levels identical to *glmU* knockdown alone (Fig. 3D). This epistatic relationship places UDP-GlcNAc upstream of the c-di-GMP regulatory node and suggests that *P. aeruginosa* monitors UDP-GlcNAc flux across envelope biosynthetic branches to fine-tune signaling outputs. Consistent with this model, *glmU* knockdown significantly reduced the c-di-GMP response to sub-MIC fosfomycin (Fig. S7). Together, these results indicate that UDP-GlcNAc synthesis is required for c-di-GMP activation in response to inhibition of both PG and LPS pathways, supporting a model in which UDP-GlcNAc utilization functions as a metabolic checkpoint for detecting cell-envelope stress and triggering c-di-GMP-dependent responses.

### Multiple PDEs contribute to the fosfomycin-induced c-di-GMP response

To determine whether c-di-GMP synthesis or degradation is affected by inhibition of cell envelope synthesis, we screened a mutant collection containing in-frame deletions of individual c-di-GMP DGCs and PDEs(23). Each mutant was transformed with the c-di-GMP reporter, and reporter activity was measured throughout growth curve with or without fosfomycin. The top panel of Fig. 4 shows c-di-GMP reporter activity in individual mutants relative to WT under untreated planktonic growth conditions, spanning exponential to early stationary phase. Under these conditions, SiaD, SadC, and MucR were the primary DGCs contributing to c-di-GMP synthesis, while RbdA, BifA, and DipA were the main PDEs responsible for degradation (Fig. 4, top). For each mutant, we compared c-di-GMP reporter activity in the presence and absence of fosfomycin and plotted the fosfomycin-induced changes relative to WT (Fig. 4, bottom). Relative to untreated controls, the *ΔdipA, ΔrocR, ΔrmcA*, Δ*bifA* mutants exhibited significantly attenuated c-di-GMP responses to fosfomycin across multiple time points, suggesting that these PDEs play key roles in mediating the fosfomycin response.

**Fig. 4.**
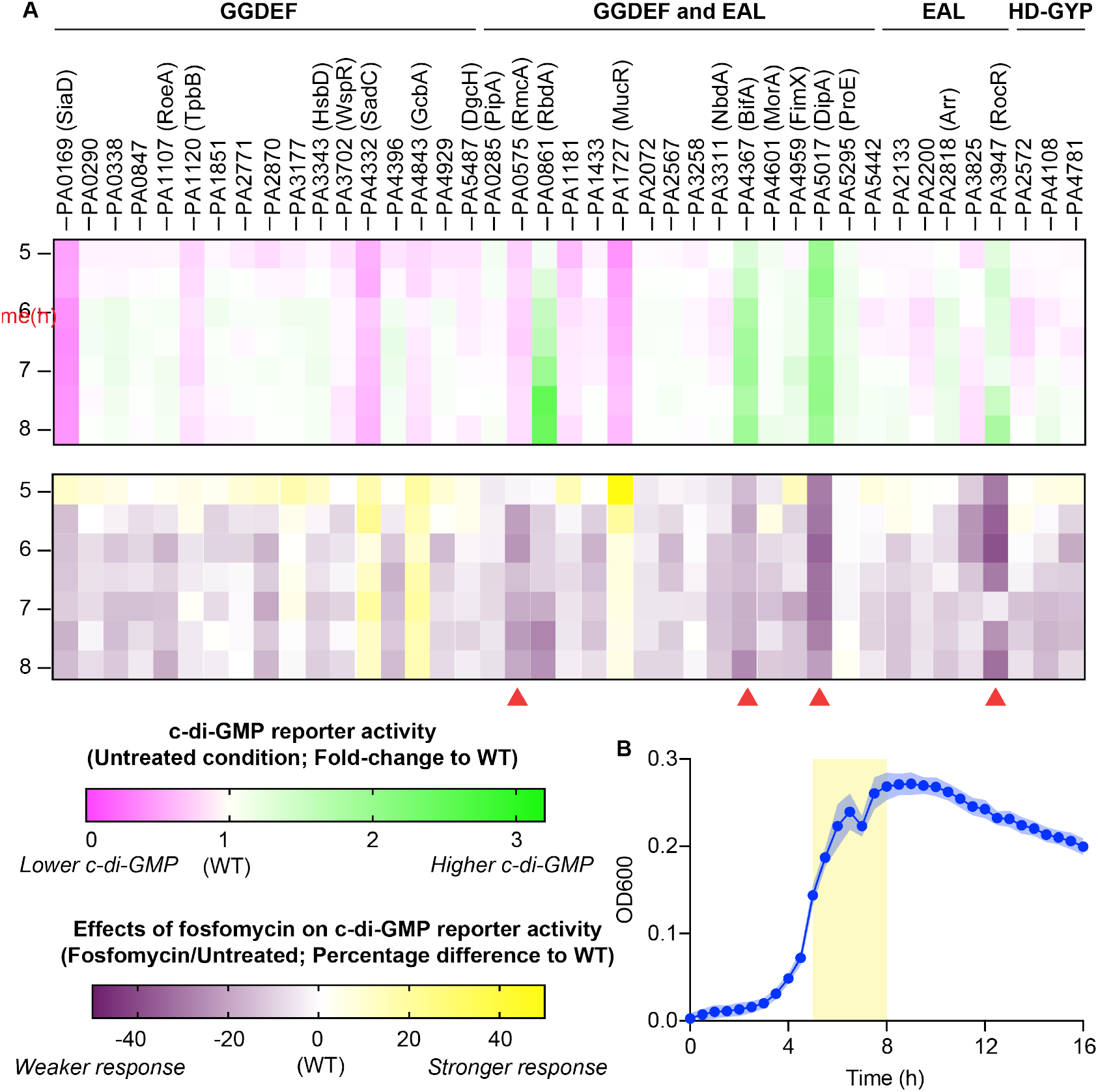
C-di-GMP reporter activity in individual c-di-GMP DGC/PDE mutants. (A) (Top) Relative c-di-GMP reporter activity compared to WT during 5-8 h of growth. Pink indicates reduced c-di-GMP reporter activity, and green indicates increased activity. WT values are normalized to 1 at each timepoint. (Bottom) Relative c-di-GMP reporter activity in response to fosfomycin compared to WT during 5-8 h of growth. The percent change in c-di-GMP reporter activity in the presence of fosfomycin was calculated for each mutant at each time point, and normalized to WT (WT = 0). Purple indicates a reduced response to fosfomycin, and yellow indicates an increased response. (B) Representative growth curve of WT. The reporter activity shown in (A) represents mid-exponential phase to early stationary phase.

However, despite attenuated responses in several PDE mutants, all mutants retained some degree of c-di-GMP reporter induction upon fosfomycin treatment (Fig. S8). These results suggest that multiple PDEs act in concert and that cell envelope stress modulates several PDEs simultaneously to elevate c-di-GMP, rather than acting through a single enzyme.

To directly visualize how c-di-GMP levels respond to cell envelope perturbation, we monitored real-time c-di-GMP dynamics in WT *P. aeruginosa* containing the cdGreen reporter, which sensitively tracks intracellular c-di-GMP fluctuations (24). After inoculation from liquid culture onto an agarose pad, c-di-GMP levels initially increased modestly during the first 30 minutes, followed by a progressive decline thereafter (Fig. 5A-B). In contrast, sub-MIC fosfomycin treatment stabilized c-di-GMP levels during this early phase, maintaining elevated levels across the population (Fig. 5A-B). Notably, cells displayed a bimodal distribution of c-di-GMP levels, indicating heterogeneous responses to envelope stress at the single-cell level (Fig. 5B). We tracked single bacteria lineages during the first 150 minutes of growth in the agarose pad (Fig. 5C, S9A-B). In the absence of fosfomycin, most lineages exhibited a homogenous reduction in c-di-GMP levels, indicating high PDE activity and low DGC activity (Fig. 5C, S9A). In the presence of fosfomycin, cells within the same lineage may respond differently, with sibling cells from the same division exhibiting distinct c-di-GMP levels, and individual cells undergoing dynamic fluctuations in which c-di-GMP levels first declined and then rose (Fig. 5C, S9B). Importantly, fosfomycin treatment did not affect individual cell growth rates (Fig. S9C), indicating that overall growth was not impaired under these conditions. These observations suggest that cell envelope stress delays c-di-GMP degradation, sustaining elevated c-di-GMP levels. However, individual cells exhibit heterogeneous regulation of PDE activity in response to cell envelope stress, leading to distinct temporal dynamics of c-di-GMP signaling.

**Fig. 5.**
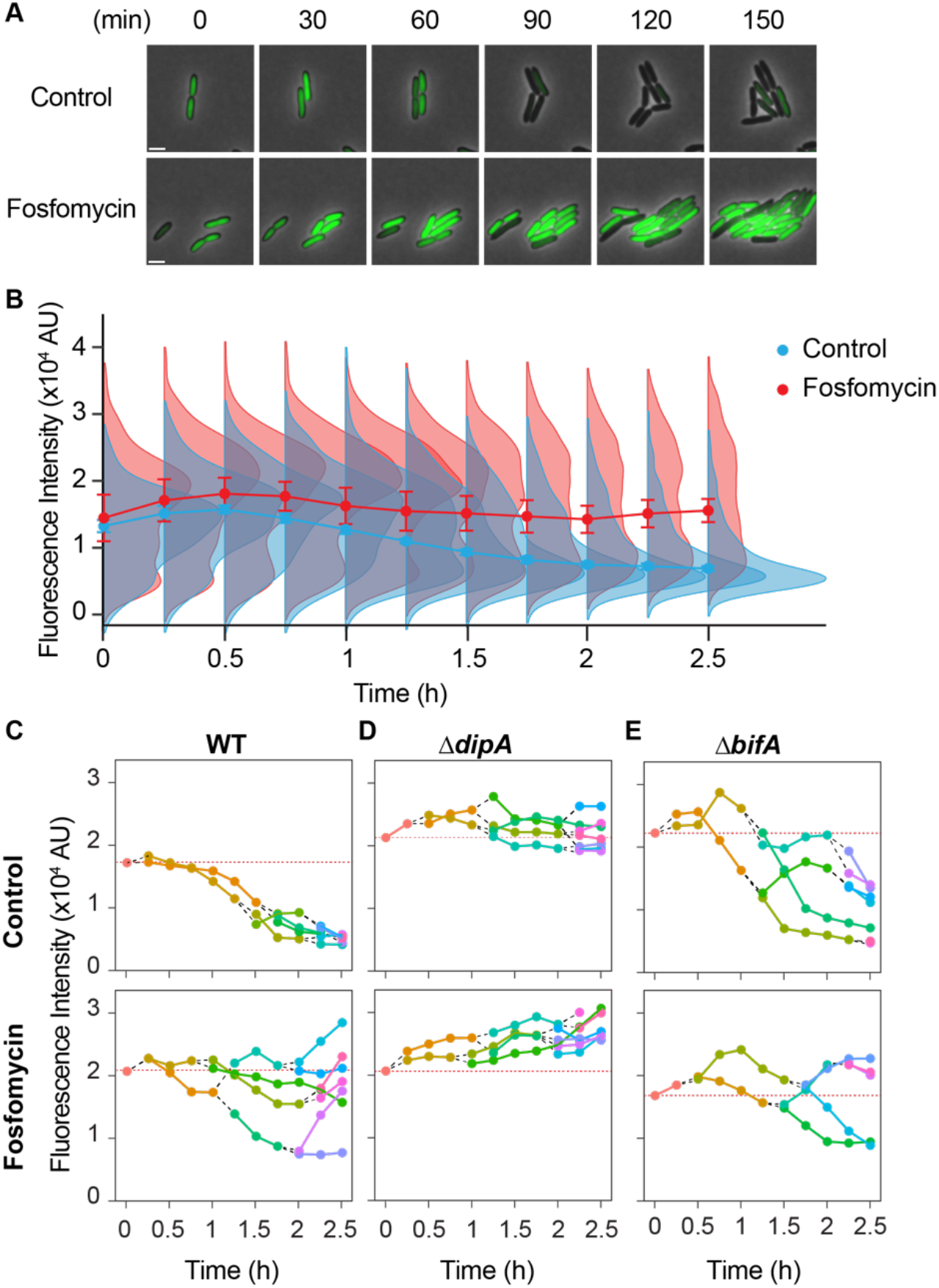
Live-cell imaging of c-di-GMP dynamics in response to fosfomycin. (A) Levels of c-di-GMP measured by cdGreen fluorescence in exponential phase bacteria inoculated onto a 1% agarose pad with or without fosfomycin. Images are representative micrographs of the same field of view. Scale bar, 2 µm. (B) Ridgeline plots of the distribution of single-cell c-di-GMP levels (cdGreen fluorescence) over time. Points with connecting lines indicate the median fluorescence at each time point. (C-E) Representative cell lineages of indicated strains in control and fosfomycin treated conditions. Points with connecting lines indicate measurement of the same cell over time. Black dash lines mark cell division events, and orange dotted lines represent cdGreen2 fluorescence at the inoculum.

Consistent with the results obtained using the *PcdrA* c-di-GMP reporter, no single PDE appears to be solely responsible for this fosfomycin-induced delay in c-di-GMP degradation. In the Δ*dipA* mutant, c-di-GMP level remained consistently high with little fluctuation without treatment, suggesting a balanced state between PDE and DGC activities (Fig. 5D). Upon fosfomycin treatment, c-di-GMP levels steadily increased over time, indicating that synthesis exceeded degradation (Fig. 5D). In the *ΔbifA, ΔrocR*, and

*ΔrmcA* mutants, untreated cells displayed heterogeneous decreases in c-di-GMP, consistent with an overall reduction in PDE activity. However, by 150 min, most cells exhibited lower c-di-GMP levels than the inoculum (Fig. 5D, S9D). In contrast, when fosfomycin was present, cells displayed a bimodal distribution, with subpopulations showing either higher or lower c-di-GMP levels compared to the inoculum (Fig. 5D, S9D). The Δ*rmcA* mutant exhibited reduced cdGreen2 fluorescence throughout the experiment, and fosfomycin treatment increased its c-di-GMP levels. Overall, live-cell imaging with the cdGreen2 reporter revealed that c-di-GMP degradation occurs over time on the agarose pad but that bacterial responses to fosfomycin are heterogeneous, leading to delayed and variable c-di-GMP degradation among individual cells.

### Fosfomycin treatment affects c-di-GMP PDE but not DGC activity

To directly assess the kinetics of c-di-GMP synthesis and degradation influenced by cell envelope stress, we optimized *in vitro* PDE and DGC assays using purified cdGreen2, cell lysates, and c-di-GMP or GTP as substrates, respectively. To validate these assays, we utilized strains overexpressing either the PDE PA2133 or the DGC PA1120. These experiments were performed in a *ΔpelΔpslΔcdrA* mutant background, which lacks the major extracellular matrix components, thereby minimizing aggregation caused by elevated c-di-GMP levels. Compared to the empty vector control (pJN105), lysates from the PA2133-overexpressing strain showed a significantly elevated rate of c-di-GMP degradation (Fig. S10A), whereas lysates from the PA1120-overexpressing strain showed a markedly higher rate of c-di-GMP synthesis (Fig. S10B). These changes in cdGreen2 fluorescence were dependent on the activity of overexpressed proteins, as the heat-inactivated cell lysate had minimal effects on cdGreen2 fluorescence.

We next prepared cell lysates from cultures grown to early stationary phase in the presence or absence of sub-MIC fosfomycin. Lysates from fosfomycin-treated cells consistently exhibited a slowed rate of c-di-GMP degradation (Fig. 6A-B). Although the absolute PDE activity varied somewhat between experiments (Fig. S11A), fosfomycin-treated lysates consistently retained approximately 70% of the PDE activity observed in untreated controls (Fig. 6B). In contrast, despite variability in c-di-GMP synthesis kinetics across experiments (Fig. S10B), fosfomycin treatment did not significantly affect the rate of c-di-GMP synthesis (Fig. 6C-D). Together with our earlier observations of delayed c-di-GMP degradation in single cells upon fosfomycin treatment, these *in vitro* assays indicate that cell envelope stress selectively inhibits PDE activity, thereby elevating c-di-GMP signaling in response to envelope perturbations.

**Fig. 6.**
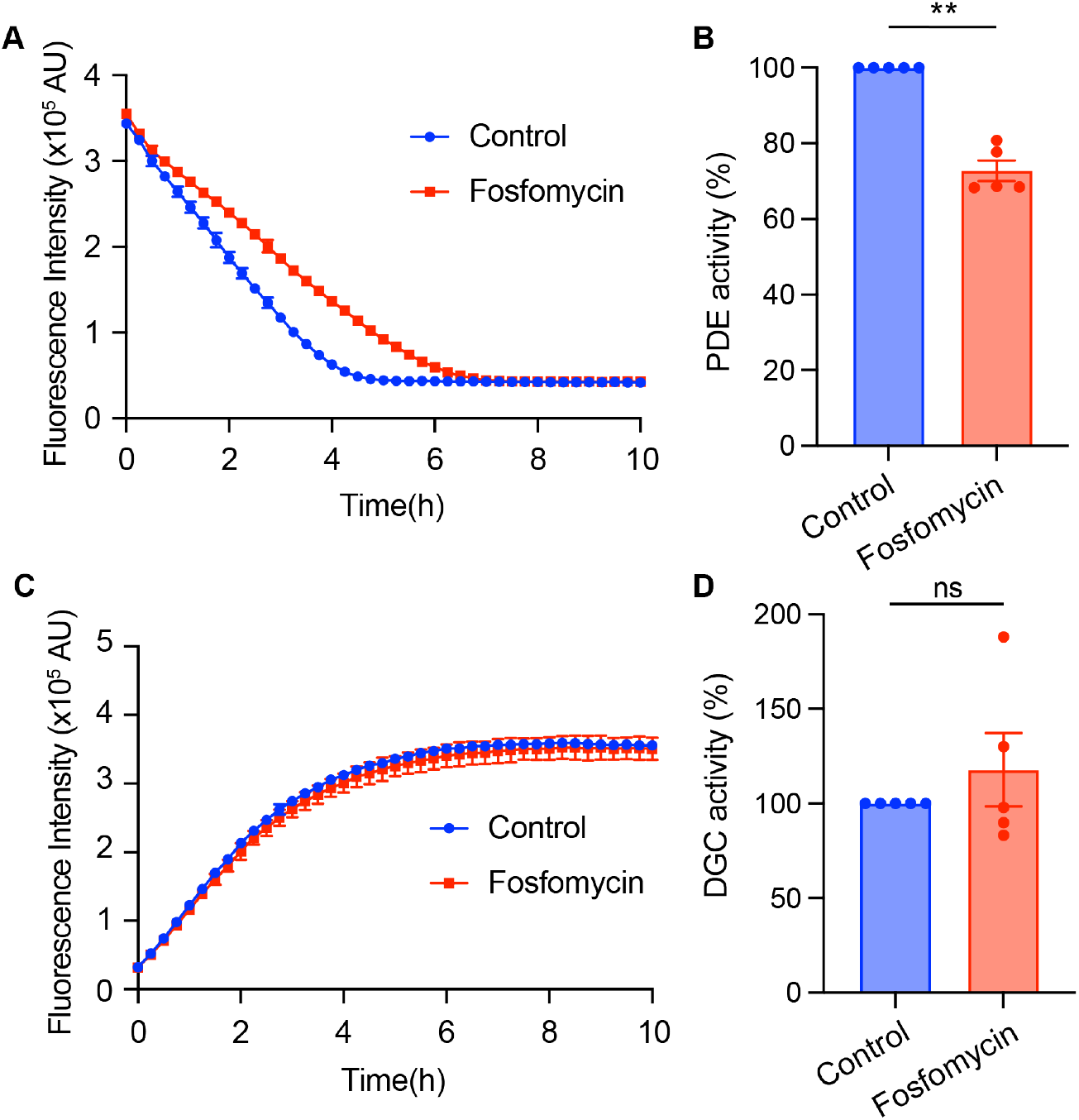
*In vitro* c-di-GMP DGC and PDE activity assay. (A) c-di-GMP degradation curves from cell lysates grown with or without 100 µg/ml fosfomycin. Data show mean ± SEM of technical replicates from a representative experiment. Results from additional biological replicates are shown in Fig. S10A. (B) Quantification of c-di-GMP PDE activity derived from degradation curves. Data are normalized to the control in each experiment and represent show mean ± SEM of independent experiments. *\*\*p* < 0.01 determined by a two-tailed t test. (C) c-di-GMP synthesis curves from cell lysates grown with or without 100 µg/ml fosfomycin. Data show mean ± SEM of technical replicates from a representative experiment. Results of additional biological replicates can be found in Fig. S10B. (D) Quantification of c-di-GMP DGC activity derived from synthesis curves. Data are normalized to the control in each experiment and represent show mean ± SEM of independent experiments. Statistics determined by a two-tailed t test.

### Acetyl-CoA, but not UDP-GlcNAc, directly regulates PDE activity

We next sought to determine if UDP-GlcNAc influences PDE activity directly. Cell lysates were pre-incubated with UDP-GlcNAc prior to initiating the PDE reaction by adding c-di-GMP and cdGreen2. Concentrations up to 1 mM of UDP-GlcNAc, as well as its epimer UDP-GalNAc, had no effect on c-di-GMP PDE activity in the lysate (Fig. 7A-B, Fig. S12A-B).

**Fig. 7.**
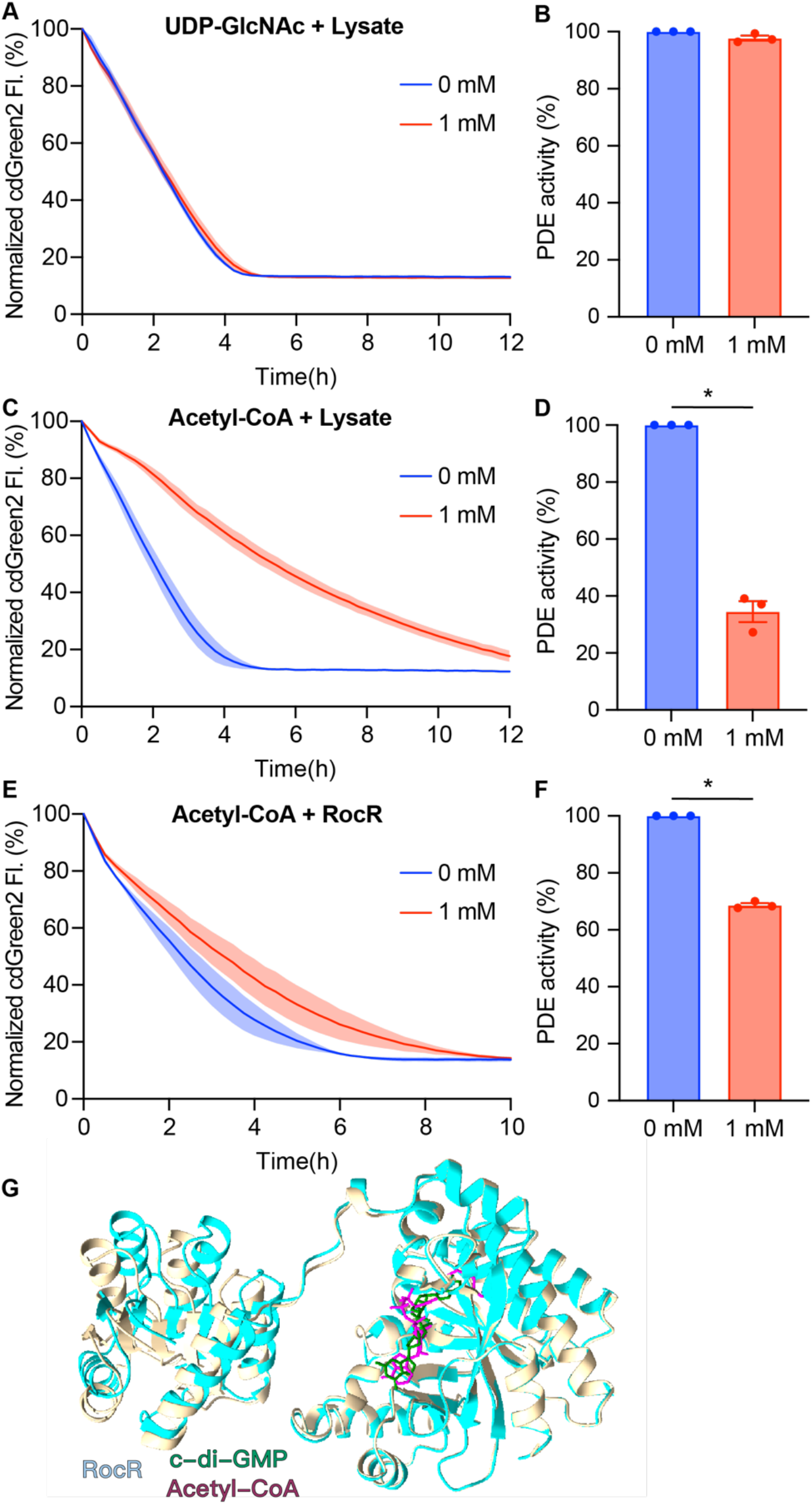
Acetyl-CoA inhibits c-di-GMP PDE activity. (A) c-di-GMP degradation curves from cell lysates with or without 1 mM UDP-GlcNAc. Data show mean ± SEM of 3 independent experiments. (B) Quantification of c-di-GMP PDE activity derived from (A). Data are normalized to the control in each experiment and represent show mean ± SEM of independent experiments. (C) c-di-GMP degradation curves from cell lysates with or without 1 mM acetyl-CoA. Date show mean ± SEM of 3 independent experiments. (D) Quantification of c-di-GMP PDE activity derived from (C). Data are normalized to the control in each experiment and represent show mean ± SEM of independent experiments. *\*\*\*\*p* < 0.0001 determined by a two-tailed t test. (E) c-di-GMP degradation curves by purified RocR with or without 1 mM acetyl-CoA. Date show mean ± SEM of 3 independent experiments. (F) Quantification of c-di-GMP PDE activity derived from (E). Data are normalized to the control in each experiment and represent show mean ± SEM of independent experiments. *\*\*\*\*p* < 0.0001 determined by a two-tailed t test. (G) Prediction of RocR structure with c-di-GMP or acetyl-CoA.

**Fig. 8.**
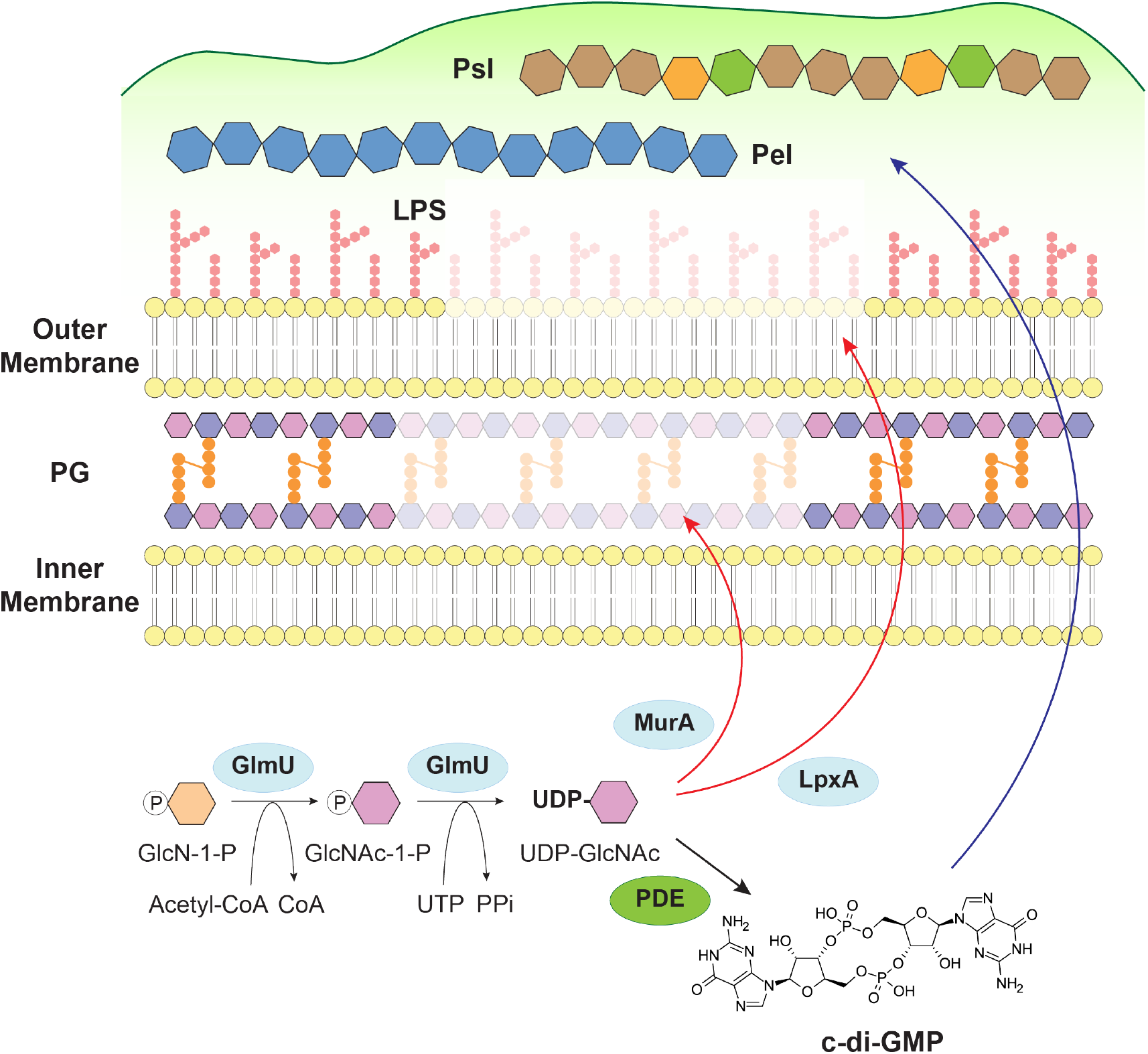
Model of bacterial coordination between cell envelope synthesis and c-di-GMP signaling. UDP-GlcNAc is synthesized from fructose-6-phosphate by the sequential action of GlmS, GlmM, and GlmU. The bifunctional enzyme GlmU consumes acetyl-CoA as a substrate in the acetyltransferase step of this pathway. The resulting UDP-GlcNAc is then channeled into multiple cell envelope biosynthetic pathways: MurA catalyzes the first committed step of peptidoglycan (PG) synthesis, while LpxA initiates lipid A synthesis as the first step of lipopolysaccharide (LPS) assembly. UDP-GlcNAc is also utilized for the synthesis of the exopolysaccharide Pel via the epimer UDP-GalNAc. When UDP-GlcNAc consumption into PG or LPS synthesis is reduced, for example, by antibiotic inhibition of MurA (fosfomycin) or CRISPRi inhibition of *lpxA*, c-di-GMP PDE activity is reduced, thereby slowing c-di-GMP degradation. The resulting increase in intracellular c-di-GMP triggers the production of exopolysaccharides Pel and Psl, which associate with the bacterial surface and form a protective extracellular matrix. This c-di-GMP-mediated response promotes biofilm formation and protects bacteria against cell envelope-targeting antimicrobials, representing an adaptive survival strategy that couples cell envelope biosynthetic status to global c-di-GMP-dependent behavior.

Given that UDP-GlcNAc does not directly modulate PDE activity, we considered whether another metabolite linked to UDP-GlcNAc biosynthesis might serve as the proximal signal. Acetyl-CoA is consumed by GlmU in the committed step of UDP-GlcNAc synthesis, acetylation of glucosamine-1-phosphate, and therefore represents a candidate metabolite whose levels could fluctuate in concert with UDP-GlcNAc demand. We found that Acetyl-CoA potently inhibited c-di-GMP PDE activity in the cell lysates (Fig. 7C-D). Furthermore, acetyl-CoA efficiently inhibited the PDE activity of purified RocR (Fig. 7E-F), suggesting a direct interaction between acetyl-CoA and RocR.

Molecular docking results indicate that acetyl-CoA likely competes for the c-di-GMP binding pocket of RocR (Fig. 7G). The predicted interaction between acetyl-CoA and RocR share several amino acids engaged in polar interactions with the RocR-c-di-GMP interaction (Fig. S13A-B). Most of these residues are conserved in EAL domains across species, implying that acetyl-CoA binding interferes with the PDE active site. The acetyl group contributes a hydrophobic moiety that is absent in c-di-GMP and extends into a groove of relatively high hydrophobicity in the EAL domain (Fig. S13C). Taken together, these results indicate that UDP-GlcNAc does not directly modulate c-di-GMP PDE activity, yet a related central metabolite, acetyl-CoA, may serve as a direct allosteric signal to inhibit PDE activity by competing for the enzyme’s active site.

## DISCUSSION

Bacteria constantly monitor and respond to cell envelope perturbations, integrating these signals into global regulatory networks including c-di-GMP signaling. However, the molecular link between cell envelope stress and c-di-GMP regulation has remained unclear. In this study, we show that sub-MIC concentrations of antibiotics targeting early PG synthesis steps increase intracellular c-di-GMP levels in *P. aeruginosa*. This effect requires UDP-GlcNAc biosynthesis, suggesting that perturbations in this precursor’s availability are sensed and relayed to c-di-GMP metabolism. Using a combination of live-cell imaging and *in vitro* enzymatic activity assays, we found that the increase in c-di-GMP elicited by cell envelope stress is not attributable to enhanced DGC activity, but rather to reduced PDE activity, thereby delaying c-di-GMP degradation. An unbiased screen of all DGC/PDE mutants revealed that no single enzyme solely accounted for the c-di-GMP increase upon cell envelope stress. Elevated c-di-GMP, in turn, provides protection against cell envelope stress (Fig. 7). A key downstream effect of c-di-GMP signaling is the synthesis of EPS. In planktonic cells, EPS can be found both secreted into the extracellular milieu as well as associated with the cell surface(21, 25). They have been shown to protect bacteria from killing by cell envelope-targeting antibiotics, such as colistin, polymyxin B, and biapenem(26–28). Notably, the exopolysaccharide Pel requires UDP-GlcNAc as a biosynthetic precursor. Increased Pel production driven by c-di-GMP signaling may consume the UDP-GlcNAc that accumulates when PG and LPS synthesis is inhibited. More broadly, activated c-di-GMP signaling promotes the formation of biofilms, a state that is notoriously resistant to diverse antimicrobial molecules and highly tolerant of hostile environments.

Our study identifies UDP-GlcNAc as a key molecule linking cell envelope biogenesis to c-di-GMP signaling. Inhibition of UDP-GlcNAc utilization in PG and LPS synthesis increased c-di-GMP signaling, whereas disruption of UDP-GlcNAc synthesis reduced c-di-GMP signaling (Fig. 2C, 3B-C). UDP-GlcNAc is a critical metabolic intermediate at the branch point for PG synthesis, LPS synthesis in Gram-negative bacteria, and wall teichoic acid (WTA) synthesis in Gram-positive bacteria. Mechanisms that balance the allocation of this molecule are therefore likely critical for bacterial survival. For example, it was reported that the LpxC enzyme, which catalyzes the committed second step of lipid A synthesis, was recently shown to interact with MurA to coordinate PG and LPS production across diverse bacterial species(29). In *Listeria monocytogenes*, defects in PG synthesis caused by *gpsB* deletion can be rescued either by *murA* overexpression or by deleting genes that divert UDP-GlcNAc into WTA synthesis(30), suggesting coordination between PG and WTA synthesis. However, how cells sense UDP-GlcNAc levels to direct flux into the appropriate pathways remains poorly understood.

Here, we demonstrate coordination between UDP-GlcNAc metabolism and c-di-GMP signaling. However, the effect of UDP-GlcNAc is likely indirect. Up to 1 mM of UDP-GlcNAc had minimal influence on c-di-GMP PDE activity. Instead, we identified acetyl-CoA, a central metabolite consumed during UDP-GlcNAc biosynthesis, as a direct inhibitor of PDE activity, potentially acting by competing for the EAL domain active site. We propose a model in which GlmU activity serves as the metabolic link between UDP-GlcNAc flux and acetyl-CoA. Under normal growth conditions, GlmU consumes acetyl-CoA to acetylate glucosamine-1-phosphate, keeping intracellular acetyl-CoA at levels insufficient to inhibit PDE activity. However, when UDP-GlcNAc consumption by PG or LPS synthesis is reduced, as occurs during fosfomycin treatment or *lpxA* knockdown, the demand on GlmU decreases, acetyl-CoA accumulates, and PDE activity is inhibited, elevating c-di-GMP. This model also accounts for the apparently paradoxical observation that *glmU* knockdown reduces rather than elevates c-di-GMP (Fig. 3) by directly impairing GlmU acetyltransferase activity, *glmU* knockdown prevents acetyl-CoA from being channeled through the UDP-GlcNAc pathway altogether, but simultaneously eliminates the flux-sensing step itself, such that perturbations downstream in PG or LPS synthesis can no longer be relayed to acetyl-CoA accumulation and PDE inhibition.

Both UDP-GlcNAc and acetyl-CoA are present at high intracellular concentrations. For exponentially growing *E. coli*, UDP-GlcNAc ranges 1-10 mM and acetyl-CoA ranges 0.2-2.0 mM(31). This contrasts with the low intracellular concentration of c-di-GMP, which ranges from nanomolar to low micromolar under standard growth conditions(32, 33). We assayed PDE activity using 1 mM UDP-GlcNAc or acetyl-CoA versus 2 µM c-di-GMP, a physiologically relevant ratio. At these concentrations, acetyl-CoA directly interacts with RocR and inhibits its PDE activity. Future work will aim to determine how UDP-GlcNAc flux influences acetyl-CoA concentration. As a substrate consumed during UDP-GlcNAc synthesis, acetyl-CoA may accumulate when the synthetic flux through this pathway is reduced. Alternatively, UDP-GlcNAc may act as a signaling molecule that reprograms central carbon metabolism. Supporting this notion, in *Bacillus subtilis*, UDP-GlcNAc is has been shown to bind YvcK (also known as GlmR), an essential protein for growth on Krebs cycle intermediates(34). The function of YvcK can be complemented by its homologue in *E. coli*, suggesting a conserved mechanism(35). Future work will be needed to determine whether a similar UDP-GlcNAc effector protein exists in *P. aeruginosa* and what its function might be. Notably, our study does not exclude the possibility that UDP-GlcNAc and acetyl-CoA serve as independent signals that converge on c-di-GMP regulation.

As a key environmental response pathway, c-di-GMP signaling is governed by a complex network. Many c-di-GMP DGCs or PDEs contain transmembrane helices or periplasmic domains that sense signals originating from the periplasm. Others contain REC domains, functioning as response regulators within two- or multi-component signal transduction systems. Allosteric regulation of c-di-GMP metabolic enzymes is also prominent. C-di-GMP can bind to the I-site of some DGCs to suppress synthesis(36, 37). PDE activity is regulated by the degradation intermediate pGpG(38, 39), and by GTP, which binds to a degenerate GGDEF domain in dual GGDEF-EAL domain PDEs(40, 41). We screened a single mutant library of *P. aeruginosa* DGCs and PDEs, and found that no single enzyme solely accounted for the c-di-GMP increase upon cell envelope stress. Instead, several PDEs, including DipA, BifA, RocR, RmcA, appeared to be influenced by fosfomycin-induced changes. These PDEs are dispersed in the chromosome, exhibit diverse domain architectures, and are reported to function in distinct environmental contexts(42–45), making coordinated transcriptional regulation of all these PDEs unlikely. We reasoned that a broader, post-translational mechanism likely influences the activity of all PDEs. We identified acetyl-CoA as a highly abundant cellular metabolite that interacts with the conserved active site of the EAL domain to inhibit PDE function. Future work will characterize acetyl-CoA-PDE interactions in more physiological contexts and identify other acetyl-CoA-like molecules that may also occupy the c-di-GMP-binding pocket of EAL domains. Our findings highlight that c-di-GMP signaling responds not only directly to environmental cues via dedicated sensory domains but also to the intracellular metabolic state through allosteric modulation of its universal catalytic domains.

The identification of acetyl-CoA as a direct inhibitor of PDE activity positions it within a broader, emerging framework in which this central metabolite serves as a signaling molecule that couples metabolic state to cellular behavior. In bacteria, intracellular acetyl-CoA concentrations fluctuate dynamically with growth phase and carbon source availability, reflecting the balance between glycolytic flux, the tricarboxylic acid cycle, and biosynthetic demands. In *Bacillus subtilis*, the AcuA acetyltransferase forms a tight regulatory complex with acetyl-CoA synthetase (AcsA) to provide feedback control of acetyl-CoA biosynthesis, illustrating how cells have evolved dedicated mechanisms to sense and respond to acetyl-CoA levels(46). Dynamic, growth-phase-dependent changes in protein acetylation state, shifting between exponential and stationary phase, have also been reported in *B. subtilis*, reinforcing the concept that acetyl-CoA fluctuations are a physiologically relevant signal tied to nutritional and growth status(47). Strikingly, the role of acetyl-CoA as a metabolic sensor is conserved in eukaryotes, where it serves as a substrate for histone acetyltransferases, directly linking nutrient availability and metabolic flux to chromatin remodeling and transcriptional regulation. Our finding that acetyl-CoA directly inhibits the EAL domain of PDEs adds a new dimension to this emerging paradigm: rather than acting solely through covalent protein modification, acetyl-CoA can also modulate protein function through direct competitive binding at a conserved catalytic site.

Given that EAL domain residues predicted to contact acetyl-CoA are broadly conserved across bacterial species, this mechanism may represent a widespread strategy by which bacteria integrate their metabolic state, specifically the demand on acetyl-CoA for cell envelope precursor biosynthesis, with c-di-GMP-dependent adaptive responses including biofilm formation.

## MATERIALS AND METHODS

### Bacterial strains and growth conditions

The bacterial strains, plasmids, and primers used in this study are listed in Tables S2-S4. *P. aeruginosa* strain PAO1 was used as the wild-type (WT) strain. *E. coli* strains were routinely cultured on LB (Lennox) agar or in LB (Lennox) broth at 37°C with shaking at 225 rpm. *P. aeruginosa* was grown on LB (Lennox) agar or in photosynthetic media(48) supplemented with 0.25% casamino acid (PM+CAA). When appropriate, antibiotics were added at the following concentrations: 30 µg/ml gentamicin or 50 µg/ml kanamycin for *E. coli*, and 100 µg/ml gentamicin for *P. aeruginosa*. Unless otherwise specified, *P. aeruginosa* were grown overnight in PM+CAA at 37°C before sub-culturing into PM+CAA to the desired growth phase for subsequent experiments.

### Generation of c-di-GMP reporter plasmids

To construct the c-di-GMP reporter, we amplified PcdrA-mGL (ASV), T2Te terminators, PrpoD, and mScarlet-I(ASV) from a previously reported tri-color reporter plasmid (XZ198) using primer pairs oXZ213/oXZ206, oXZ225/oXZ229, oXZ216/oXZ21, oXZ22/oXZ230, respectively. The resulting four fragments were assembled into the SacI/NcoI-digested pBBR1MCS5 vector using HiFi assembly, resulting in XZ197 reporter. To generate the c-di-GMP reporter containing a constitutively expressed sgRNA, P*speI* along with the gRNA scaffold was amplified from pPSV37(49) using primers MO172/MO173 and inserted into the XmaI-digested XZ197 plasmid, resulting in the empty vector control plasmid. From this empty vector, sgRNAs targeting different genes were synthesized as primers, and paired with the universal reverse primer (oXZ434) to amplify the entire reporter plasmid. sgRNAs were designed to be 25 nt in length, on the non-template strand, adjacent to a 5’-NGG PAM sequence, and located close to the start of the transcript. The PCR products were circularized by HiFi assembly. To introduce a second CRIPSRi target downstream of the CRISPRi-c-di-GMP reporter, the single-sgRNA plasmid targeting *glmU* or the empty vector was PCR amplified using primers oXZ581/oXZ582 to linearize. A second copy of P*speI*-gRNA scaffold-T7 terminator targeting *murB, lpxA, glmU*, or an empty vector was amplified using primers oXZ579/oXZ580, and HiFi assembled into the linearized plasmid.

### Plate reader-based measurement of bacterial growth and c-di-GMP reporter activity

To assess bacterial growth and the c-di-GMP reporter activity in WT, the overnight culture was diluted 1:100 into 100 µl of PM+CAA+100 µg/ml gentamicin in a clear-bottom black 96-well plate. For the initial stressor screen, the medium contained 1:2 serial dilutions of test molecules. For subsequent experiments, the medium was supplemented with specific molecules at defined concentrations, or with 0.2% arabinose to induce dCas9 expression. Plates were sealed with a Breathe-Easy membrane (Sigma-Aldrich) and incubated in a ClarioStar Plus plate reader (BMG Labtech) at 37°C with shaking at 200 rpm. The growth (OD600), c-di-GMP reporter fluorescence (mGL; Ex490/10, Em530/20), and constitutive reporter fluorescence (mScarlet-I; Ex570/15, Em613/20) were recorded every 30 min for 18 hours.

### Microscopy-based measurement of c-di-GMP reporter activity

To evaluate single-cell c-di-GMP reporter activity, WT carrying the c-di-GMP reporter with either an empty gRNA scaffold or an sgRNA targeting *murB* were grown with or without 0.1% arabinose until the empty vector strain each an OD600 of ∼0.9. Cells were fixed with 2.5% paraformaldehyde (PFA) for 10 min at 37°C and imaged using a Nikon Ti2 microscope. GFP (Ex466/40, Em525/50) filter was used for the c-di-GMP reporter, and TRITC (Em554/23, Em609/54) filter for the constitutive reporter.

### Quantification of c-di-GMP

Quantification of c-di-GMP by liquid chromatography-mass spectrometry (LC-MS) was performed as previously described(50). Briefly, bacterial cultures were harvested at early-stationary phase (OD600 ∼0.9). Cells were collected by centrifugation at 4,000xg, 10min at 4°C, and proteins were precipitated by incubation with perchloric acid (PCA) on ice for 1 h. An internal control of 2-chloro AMP was included in each sample, and the supernatant was neutralized by potassium bicarbonate. LC-MS was performed using an Acuity UPLC with a Synergi 4 µ Hydro RP 80A column and a C18 Guard Cartridge (Phenomenex) on a Xevo TQ-S mass spectrometer (Waters). The mass transition monitored were m/z 691 > 152 for c-di-GMP and 382 > 170 for 2-chloro-AMP. For protein quantification, PCA-precipitated pellets were neutralized with NaOH and solubilized by boiling for 60 min in SDS sample buffer. Protein concentration was measured by Pierce 660nm protein assay with the ionic detergent compatibility reagent (ThermoFisher).

### Time-lapse microscopy

To monitor c-di-GMP dynamics using the cdGreen2 reporter, bacteria were cultured to mid-exponential phase (OD600 ∼0.4) in PM+CAA supplemented with gentamicin. A 1% agarose pad was prepared in 1 ml PM+CAA with or without 100 µg/ml fosfomcyin, and solidified between two coverslips at room temperature for 45 min. Bacterial culture was inoculated at the center of the agarose pad, and the pad was placed on a glass bottom dish (MatTek P35G-1.5-14-C). Water was added to the side of the dish to reduce evaporation. The same field of view was imaged with every 15 minutes at 30°C using a Nikon Ti2 microscope with the GFP (Ex466/40, Em525/50) filter.

Image analysis was carried out using the OmniSegger pipeline(51). The Omnipose(52) segmentation was performed using the bact_phase_affinity pretrained model, mask_threshold of 1, and flow_threshold of 0. The distribution of mean fluorescence per cell was visualized as ridgeline plots, and the median fluorescence across all cells were calculated and plotted as a line plot in R. The representative lineages were selected based on having a single isolated bacterium as the ancestor of the lineage and at least two cell divisions, and lineage relationships were curated manually.

### cdGreen2 purification

cdGreen2 protein was purified as previously described(24) with modifications. *E. coli* BL21 containing pET28-cdGreen2 was cultured in 1.5 L terrific broth to an OD600 of ∼0.8, and expression was induced with 0.1mM IPTG overnight. Cells were pelleted, washed, and lysed in lysis buffer (50mM Tris-HCl, pH 8.0, 250mM NaCl, 20mM imidazole) containing 1mg/ml lysozyme using a sonicator. Cell debris was removed by centrifugation, and the supernatant was filtered through a 0.45µm PVDF syringe filter before incubation with Ni-NTA resin (Qiagen) for 2 h at 4°C. The resin was centrifuged to remove the unbound fraction and re-suspended with the lysis buffer to load into a 1.5 × 15 cm column. The resin was washed with 50mM Tris-HCl, pH 8.0, 250mM NaCl, 40mM imidazole until the flow-through contained minimal protein, and cdGreen2 was eluted from with 50mM Tris-HCl, pH 8.0, 250mM NaCl, 300mM imidazole. The eluate was further purified by size-exclusion chromatography on a Superdex S200 column (Cytiva) in the sensor buffer (25mM Tris-HCl pH 7.5, 150mM NaCl, 10mM MgCl2, 10mM KCl, 5mM β-mercaptoethanol). Protein concentration was calculated by absorbance at 280 nm, yielding a 50 µM stock.

### *In vitro* c-di-GMP PDE and DGC assay

Measurement of c-di-GMP PDE and DGC activity in cell lysates using cdGreen2 was adapted from a previously described assay with purified proteins(24). For the control experiment (pJN105/pJN2133/pJN1120), bacterial cultures were grown with 0.2% arabinose until mid-exponential phase (OD600 ∼0.5). Cells were pelleted and lysed by sonication in PBS with 10 mM MgCl_2_. Cell debris were removed by centrifuging at 4,000 xg, 10min and filtered through a 0.22 µm PDVF syringe filter. The resulting lysates had protein concentrations ranging 0.6-0.8 mg/ml, measured by a Pierce BCA protein assay (ThermoFisher). For the assay, substrates (2 µM c-di-GMP for PDE activity, or 2mM GTP for DGC activity) were pre-equilibrated with 200 nM purified cdGreen in 100 µl PBS+10mM MgCl_2_ in a clear-bottom black 96-well plate for at least 1 h at 30°C. Reactions were initiated by adding 10 µl of cell lysate (normalized to 0.6 mg/ml) to the cdGreen/substrate mixture. Fluoresence (Ex488/405, Em530) was measured every 15 min at 30°C using a ClarioStar Plus plate reader.

To evaluate the effects of fosfomycin on PDE/DGC activity, WT bacteria were grown to early-stationary phase (OD600 ∼0.9) with or without 100 µg/ml fosfomycin. Cell lysates were prepared as above, then concentrated to at least 10 mg/ml using an Amicon Ultra-15 centrifugal filter unit (10 KDa MWCO, Millipore). The lysates were normalized to 1.6 mg/ml, and 10 µl of each sample was added to the cdGreen2/substrate mixture to initiate the reaction and measurements.

### Statistical analysis

All statistical analyses were performed in Graphpad Prism 10. Unless otherwise mentioned, each dot represents an independent experiment.

## ACKNOWLEDGEMENT

We thank the members of the Parsek laboratory for insightful discussions. We thank Dr. Urs Jenal for the gift of cdGreen reporter and purification plasmid. We thank Dr. Alain Filloux for the gift of PAO1 collection of individual DGC/PDE mutants.We thank Dr. Alexander Meeske and members of the Parsek Laboratory for scientific discussions. We thank Dr. Sora Kim for assistance in protein purification. We thank Dr. Teresa Lo and Dr. Kevin Cutler for assistance in OmniSegger analysis. This study was supported by the National Institute of Health R01AI143916 and R01AI077628 to M.R.P and K99GM160696 to X.Z. X.Z. is a Cystic Fibrosis Foundation Awardee of the Life Sciences Research Foundation.

## REFERENCES

1. U. Jenal, A. Reinders, C. Lori, Cyclic di-GMP: second messenger extraordinaire. Nat. Rev. Microbiol. 15, 271–284 (2017).

2. L. Hall-Stoodley, J. W. Costerton, P. Stoodley, Bacterial biofilms: from the natural environment to infectious diseases. Nat. Rev. Microbiol. 2, 95–108 (2004).

3. T.-F. C. Mah, G. A. O’Toole, Mechanisms of biofilm resistance to antimicrobial agents. Trends Microbiol. 9, 34–39 (2001).

4. H. Kulesekara, et al., Analysis of *Pseudomonas aeruginosa* diguanylate cyclases and phosphodiesterases reveals a role for bis-(3′-5′)-cyclic-GMP in virulence. Proceedings of the National Academy of Sciences 103, 2839–2844 (2006).

5. T. J. Silhavy, D. Kahne, S. Walker, The Bacterial Cell Envelope. Cold Spring Harb. Perspect. Biol. 2, a000414–a000414 (2010).

6. J. Vergalli, et al., Porins and small-molecule translocation across the outer membrane of Gram-negative bacteria. Nat. Rev. Microbiol. 18, 164–176 (2020).

7. K. A. Kline, S. Fälker, S. Dahlberg, S. Normark, B. Henriques-Normark, Bacterial Adhesins in Host-Microbe Interactions. Cell Host Microbe 5, 580–592 (2009).

8. K. Lewis, The Science of Antibiotic Discovery. Cell 181, 29–45 (2020).

9. A. M. Mitchell, T. J. Silhavy, Envelope stress responses: balancing damage repair and toxicity. Nat. Rev. Microbiol. 17, 417–428 (2019).

10. G. Rowley, M. Spector, J. Kormanec, M. Roberts, Pushing the envelope: extracytoplasmic stress responses in bacterial pathogens. Nat. Rev. Microbiol. 4, 383–394 (2006).

11. M. Wehland, F. Bernhard, The RcsAB Box. Journal of Biological Chemistry 275, 7013–7020 (2000).

12. C. D. Hershberger, R. W. Ye, M. R. Parsek, Z. D. Xie, A. M. Chakrabarty, The algT (algU) gene of Pseudomonas aeruginosa, a key regulator involved in alginate biosynthesis, encodes an alternative sigma factor (sigma E). Proceedings of the National Academy of Sciences 92, 7941–7945 (1995).

13. T. Cho, K. Pick, T. L. Raivio, Bacterial envelope stress responses: Essential adaptors and attractive targets. Biochim. Biophys. Acta Mol. Cell Res. 1870 (2023).

14. J. Klebensberger, O. Rui, E. Fritz, B. Schink, B. Philipp, Cell aggregation of Pseudomonas aeruginosa strain PAO1 as an energy-dependent stress response during growth with sodium dodecyl sulfate. Arch. Microbiol. 185, 417–427 (2006).

15. A. I. Chen, et al., Candida albicans Ethanol Stimulates Pseudomonas aeruginosa WspR-Controlled Biofilm Formation as Part of a Cyclic Relationship Involving Phenazines. PLoS Pathog. 10, e1004480 (2014).

16. J. Klebensberger, A. Birkenmaier, R. Geffers, S. Kjelleberg, B. Philipp, SiaA and SiaD are essential for inducing autoaggregation as a specific response to detergent stress in *Pseudomonas aeruginosa*. Environ. Microbiol. 11, 3073–3086 (2009).

17. K. A. Lewis, et al., Ethanol Decreases Pseudomonas aeruginosa Flagellar Motility through the Regulation of Flagellar Stators. J. Bacteriol. 201 (2019).

18. L. O’Neal, et al., The Wsp system of *Pseudomonas aeruginosa* links surface sensing and cell envelope stress. Proceedings of the National Academy of Sciences 119 (2022).

19. V. Soni, E. H. Rosenn, R. Venkataraman, Insights into the central role of N-acetyl-glucosamine-1-phosphate uridyltransferase (GlmU) in peptidoglycan metabolism and its potential as a therapeutic target. Biochemical Journal 480, 1147–1164 (2023).

20. J. D. King, D. Kocíncová, E. L. Westman, J. S. Lam, Review: Lipopolysaccharide biosynthesis in *Pseudomonas aeruginosa*. Innate Immun. 15, 261–312 (2009).

21. L. K. Jennings, et al., Pel is a cationic exopolysaccharide that cross-links extracellular DNA in the *Pseudomonas aeruginosa* biofilm matrix. Proceedings of the National Academy of Sciences 112, 11353–11358 (2015).

22. L. S. Marmont, et al., PelX is a UDP-N-acetylglucosamine C4-epimerase involved in Pel polysaccharide-dependent biofilm formation. Journal of Biological Chemistry 295, 11949–11962 (2020).

23. K. Eilers, et al., Phenotypic and integrated analysis of a comprehensive Pseudomonas aeruginosa PAO1 library of mutants lacking cyclic-di-GMP-related genes. Front. Microbiol. 13, 949597 (2022).

24. A. Kaczmarczyk, et al., A genetically encoded biosensor to monitor dynamic changes of c-di-GMP with high temporal resolution. Nat. Commun. 15, 3920 (2024).

25. J. E. Dreifus, et al., The Sia System and c-di-GMP Play a Crucial Role in Controlling Cell-Association of Psl in Planktonic P. aeruginosa. J. Bacteriol. 204 (2022).

26. K. Murakami, et al., Role of psl Genes in Antibiotic Tolerance of Adherent Pseudomonas aeruginosa. Antimicrob. Agents Chemother. 61 (2017).

27. N. Billings, et al., The extracellular matrix Component Psl provides fast-acting antibiotic defense in Pseudomonas aeruginosa biofilms. PLoS Pathog. 9, e1003526 (2013).

28. B. Yang, et al., Pseudomonas aeruginosa Oligoribonuclease Controls Tolerance to Polymyxin B by Regulating Pel Exopolysaccharide Production. Antimicrob. Agents Chemother. 66, e0207221 (2022).

29. K. R. Hummels, et al., Coordination of bacterial cell wall and outer membrane biosynthesis. Nature 615, 300–304 (2023).

30. J. Rismondo, J. K. Bender, S. Halbedel, Suppressor mutations linking GpsB with the first committed step of peptidoglycan biosynthesis in listeria monocytogenes. J. Bacteriol. 199 (2017).

31. B. D. Bennett, et al., Absolute metabolite concentrations and implied enzyme active site occupancy in Escherichia coli. Nat. Chem. Biol. 5, 593–599 (2009).

32. M. Christen, et al., Asymmetrical distribution of the second messenger c-di-GMP upon bacterial cell division. Science (1979). 328, 1295–1297 (2010).

33. B. R. Kulasekara, et al., c-di-GMP heterogeneity is generated by the chemotaxis machinery to regulate flagellar motility. Elife 2 (2013).

34. E. Foulquier, A. Galinier, YvcK, a protein required for cell wall integrity and optimal carbon source utilization, binds uridine diphosphate-sugars. Sci. Rep. 7 (2017).

35. B. Görke, E. Foulquier, A. Galinier, YvcK of Bacillus subtilis is required for a normal cell shape and for growth on Krebs cycle intermediates and substrates of the pentose phosphate pathway. Microbiology (N. Y). 151, 3777–3791 (2005).

36. C. Chan, et al., “Structural basis of activity and allosteric control of diguanylate cyclase” (2004).

37. B. Christen, et al., Allosteric Control of Cyclic di-GMP Signaling. Journal of Biological Chemistry 281, 32015–32024 (2006).

38. D. Cohen, et al., Oligoribonuclease is a central feature of cyclic diguanylate signaling in Pseudomonas aeruginosa. Proc. Natl. Acad. Sci. U. S. A. 112, 11359–11364 (2015).

39. M. W. Orr, et al., Oligoribonuclease is the primary degradative enzyme for pGpG in Pseudomonas aeruginosa that is required for cyclic-di-GMP turnover. Proc. Natl. Acad. Sci. U. S. A. 112, E5048–E5057 (2015).

40. B. Christen, et al., Allosteric control of cyclic di-GMP signaling. Journal of Biological Chemistry 281, 32015–32024 (2006).

41. J. C. Van Loon, et al., Binding of GTP to BifA is required for the production of Pel-dependent biofilms in Pseudomonas aeruginosa. J. Bacteriol. 206 (2024).

42. A. B. Roy, O. E. Petrova, K. Sauer, The phosphodiesterase DipA (PA5017) is essential for Pseudomonas aeruginosa biofilm dispersion. J. Bacteriol. 194, 2904–2915 (2012).

43. S. L. Kuchma, et al., BifA, a cyclic-di-GMP phosphodiesterase, inversely regulates biofilm formation and swarming motility by Pseudomonas aeruginosa PA14 in Journal of Bacteriology, (2007), pp. 8165–8178.

44. H. D. Kulasekara, et al., A novel two-component system controls the expression of Pseudomonas aeruginosa fimbrial cup genes. Mol. Microbiol. 55, 368–380 (2005).

45. S. Katharios-Lanwermeyer, G. B. Whitfield, P. L. Howell, G. A. O’Toole, Pseudomonas aeruginosa Uses c-di-GMP Phosphodiesterases RmcA and MorA To Regulate Biofilm Maintenance. mBio 12, 1–19 (2021).

46. S. Suzuki, N. Kondo, M. Yoshida, M. Nishiyama, S. Kosono, Dynamic changes in lysine acetylation and succinylation of the elongation factor Tu in bacillus subtilis. Microbiology (United Kingdom) 165, 65–77 (2019).

47. L. Zheng, et al., Regulation of acetyl-CoA biosynthesis via an intertwined acetyl-CoA synthetase/acetyltransferase complex. Nature Communications 16 (2025).

48. M. K. Kim, C. S. Harwood, Regulation of benzoate-CoA ligase in Rhodopseudomonas palustris. FEMS Microbiol. Lett. 83, 199–203 (1991).

49. A.-S. Stolle, B. T. Meader, J. Toska, J. J. Mekalanos, Endogenous membrane stress induces T6SS activity in *Pseudomonas aeruginosa*. Proceedings of the National Academy of Sciences 118 (2021).

50. Y. Irie, et al., Self-produced exopolysaccharide is a signal that stimulates biofilm formation in *Pseudomonas aeruginosa*. Proceedings of the National Academy of Sciences 109, 20632–20636 (2012).

51. T. W. Lo, K. J. Cutler, H. J. Choi, P. A. Wiggins, OmniSegger: A time-lapse image analysis pipeline for bacterial cells. PLoS Comput. Biol. 21 (2025).

52. K. J. Cutler, et al., Omnipose: a high-precision morphology-independent solution for bacterial cell segmentation. Nat. Methods 19, 1438–1448 (2022).

