## Supplementary information for "A metabolic checkpoint coordinates bacterial cell envelope biosynthesis and c-di-GMP signaling"

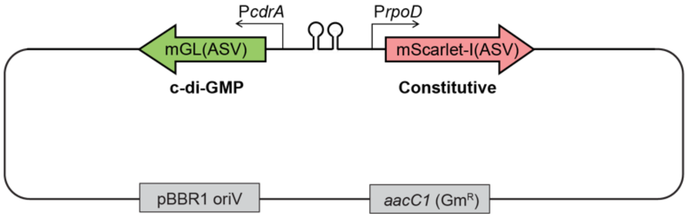


**Supplementary Fig. 1. Design of the c-di-GMP reporter plasmid.** Schematic of the c-di-GMP reporter plasmid used in this study. All fluorescent proteins were tagged with a AANDENYAASV peptide (ASV) to reduce the half-lives. The activity of the *cdrA* activity reflects c-di-GMP signaling, whereas the activity of the *rpoD* promoter serves as a constitutive control.

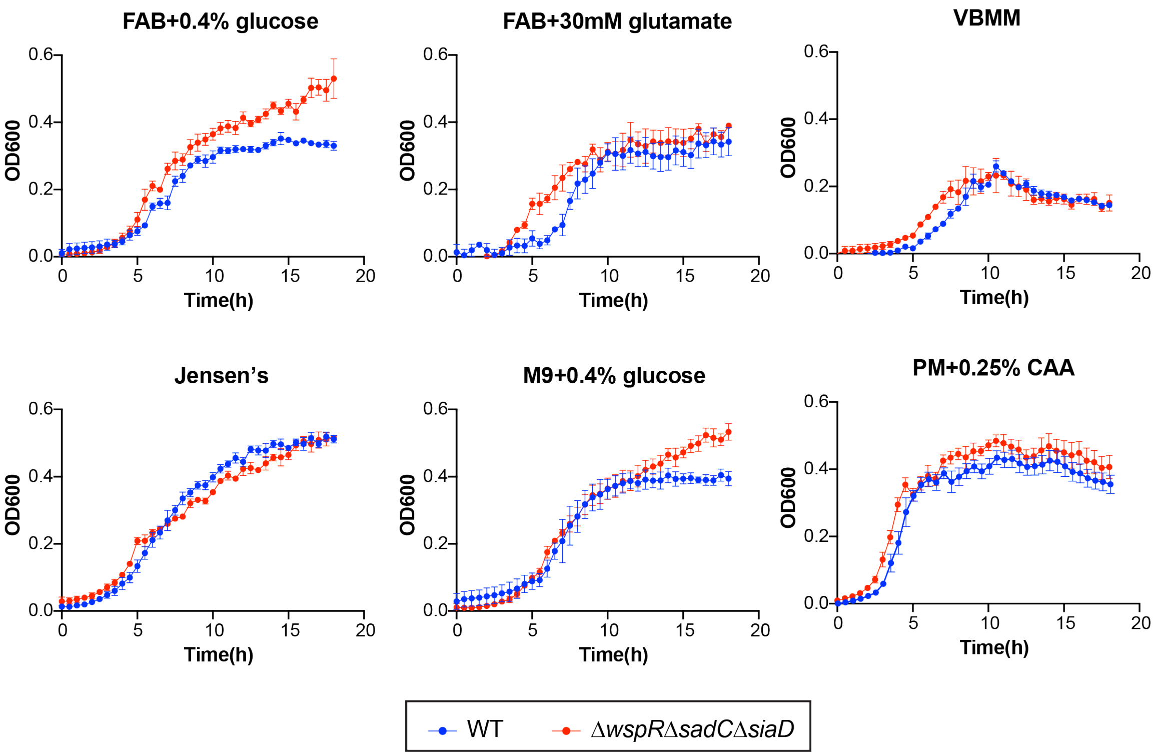


**Supplementary Fig. 2. Screening of defined media for reliable measurement of c-di-GMP reporter activity using a plate reader.** Growth curves of wild-type (WT) and a Δ*wspR*Δ*sadC*Δ*siaD* triple c-di-GMP cyclase mutant in the indicated growth media. Growth in PM+0.25% CAA showed shortest lag phase and the highest saturation OD600.

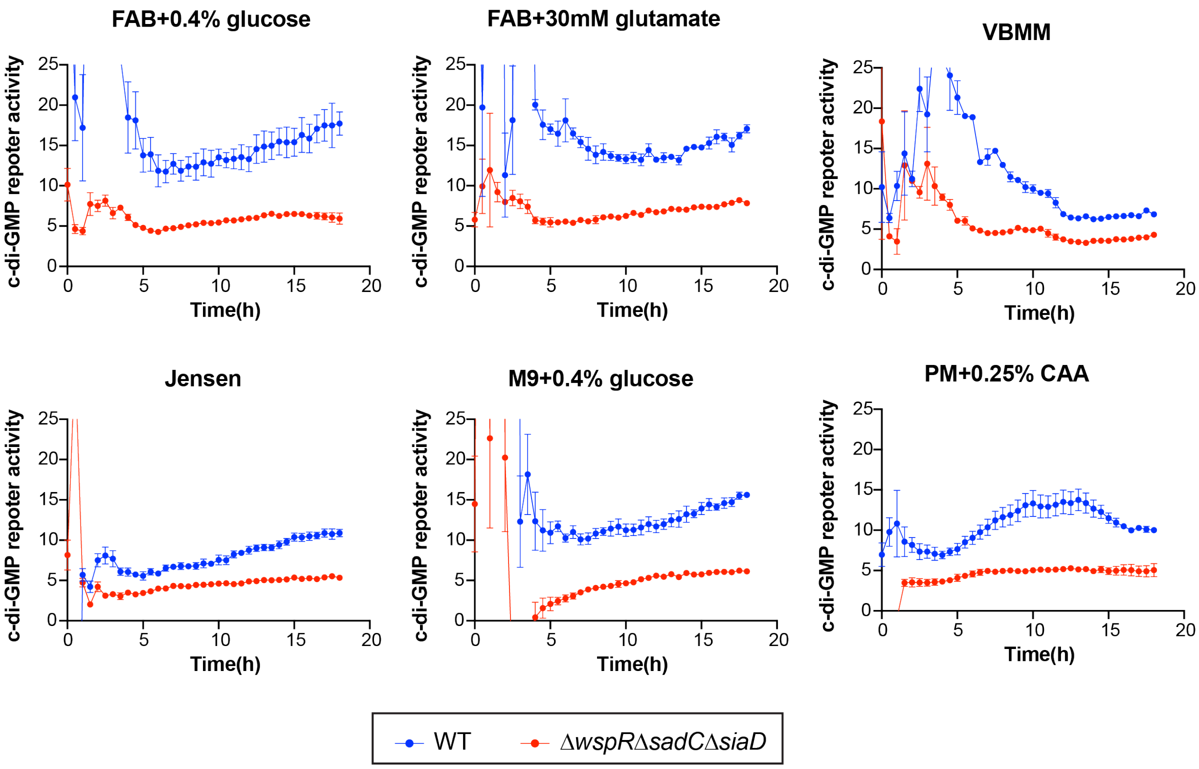


**Supplementary Fig. 3. c-di-GMP reporter activity in various defined media.** C-di-GMP reporter activity measured throughout the growth curve experiment comparing WT and the Δ*wspR*Δ*sadC*Δ*siaD* triple c-di-GMP cyclase mutant. All defined media showed a significant difference in reporter activity between WT and the triple mutant.

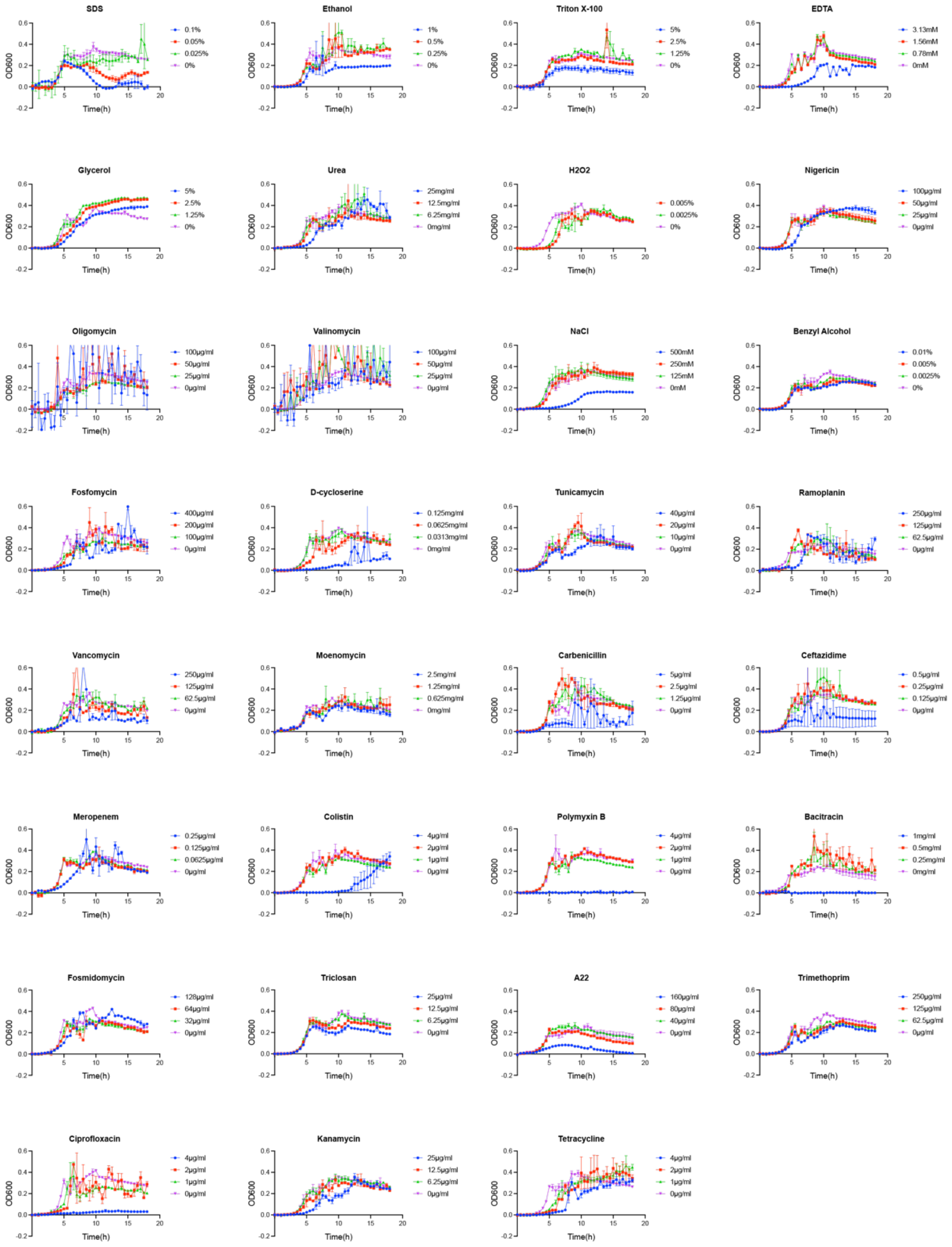


**Supplementary Fig. 4. Growth curves of *P. aeruginosa* in the panel of molecules used in the screen.** Representative OD600 measurements of WT with the indicated molecules at 1x MIC, 1/2x MIC, and 1/4x MIC concentrations. Each experiment includes an untreated control. Notably, we also compared the constitutive P*rpoD* driven mScarlet-I fluorescence to confirm the MIC for each molecule.

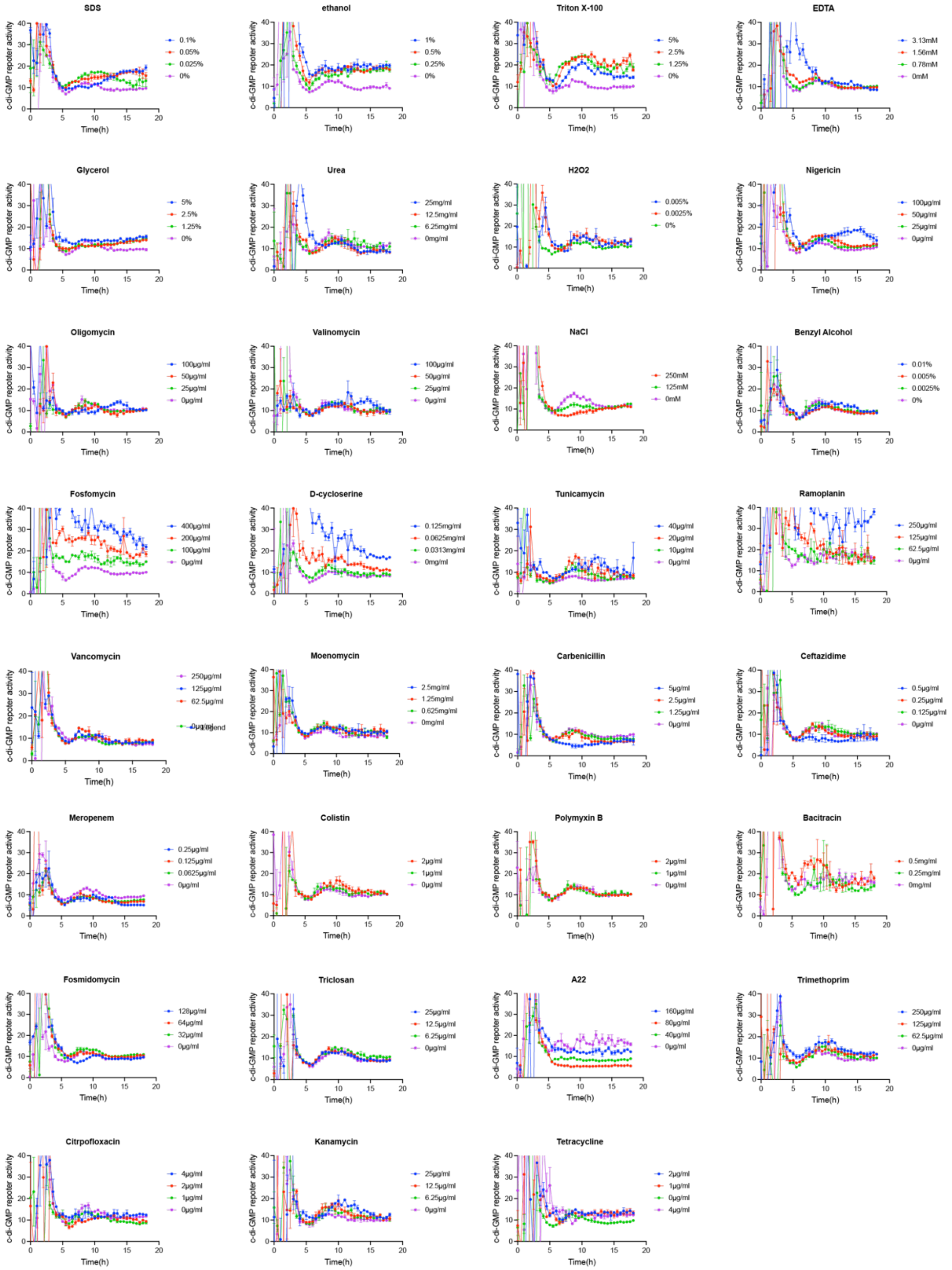


**Supplementary Fig. 5. c-di-GMP reporter activity in response to screened molecules.** Representative time-course curves of c-di-GMP reporter activity in WT grown with 1x, 1/2x, and 1/4x MIC of the indicated molecules. Each experiment includes an untreated control.

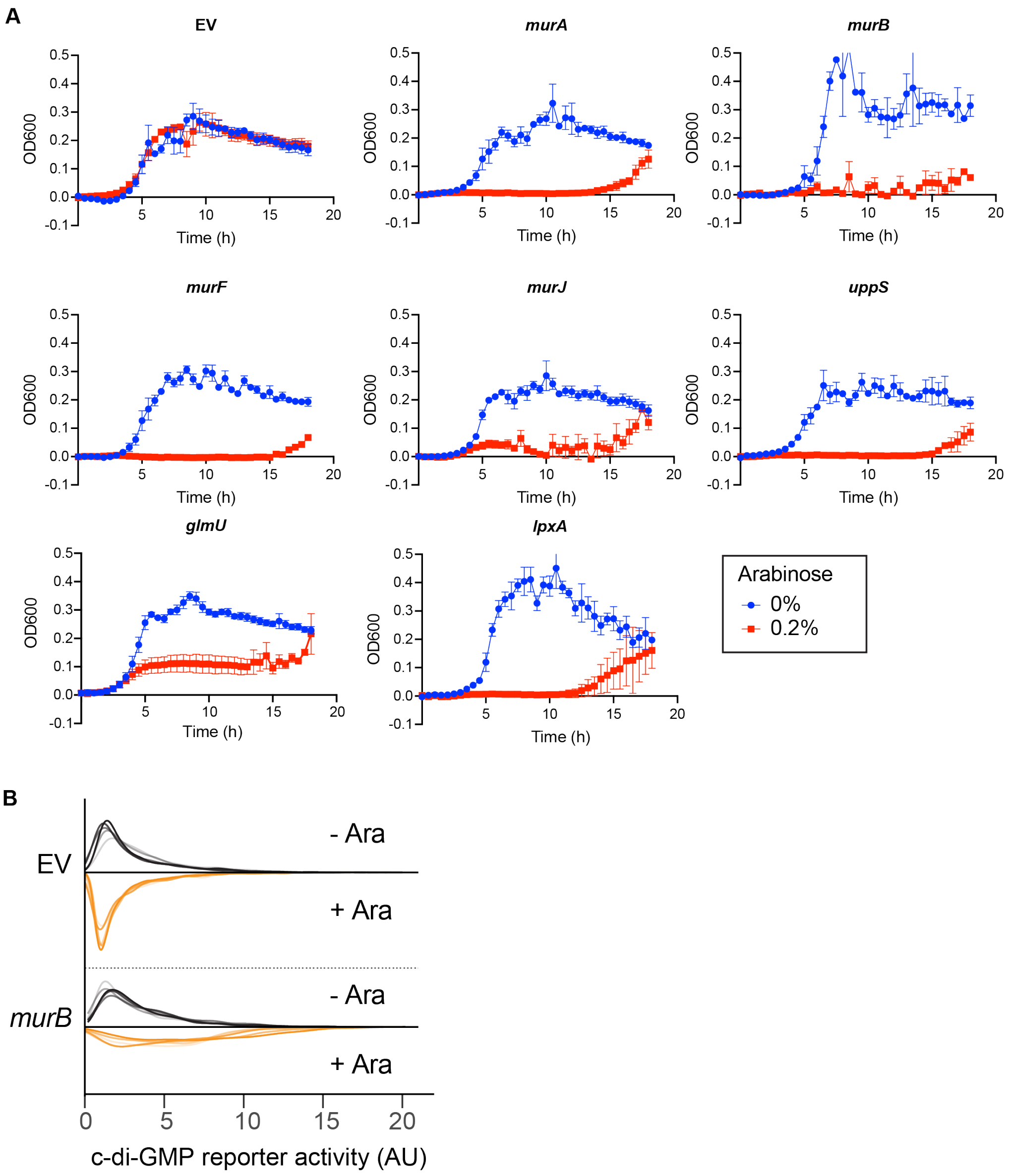


**Supplementary Fig. 6. Validation of sgRNA design and knockdown efficiency.** (A) Growth curves of CRISPRi strain expressing sgRNA targeting indicated essential genes. Effective knockdown is indicated by growth defects upon dCas9 induction with arabinose. Variability in sgRNA-target affinity likely results in differing knockdown efficiencies. (B) Single-cell distribution of c-di-GMP reporter activity in the empty vector (EV) control or *murB* knockdown strains. Arabinose has no effect on EV. Without arabinose, a small subset of *murB* knockdown cells exhibit elevated c-di-GMP reporter activity, indicating leaky dCas9 expression and the *murB* sgRNA has high efficiency. Upon arabinose induction, c-di-GMP activation is heterogeneous across the population.

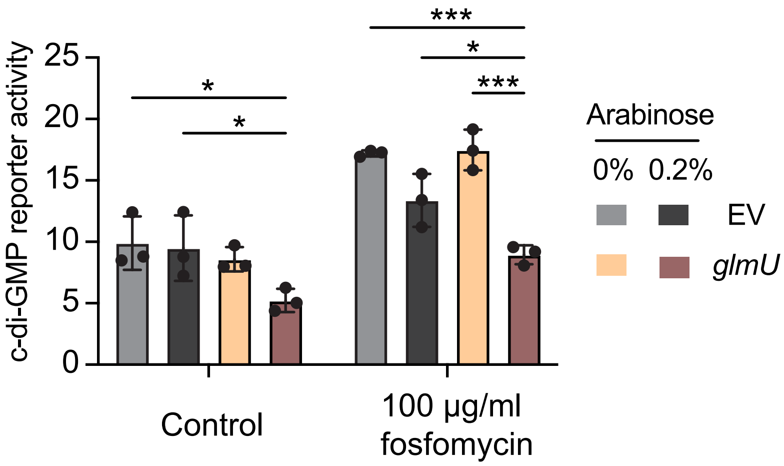


**Supplementary Fig. 7. Combined effects of *glmU* knockdown and fosfomycin.** c-di-GMP reporter activity in response to 100 µg/ml fosfomycin in strains containing an EV or CRISPRi knockdown targeting *glmU*. Arabinose induced *glmU* knockdown. *$p < 0.05$, **$p < 0.01$, ***$p < 0.001$, determined by two-way ANOVA with Sidak's multiple comparisons test.

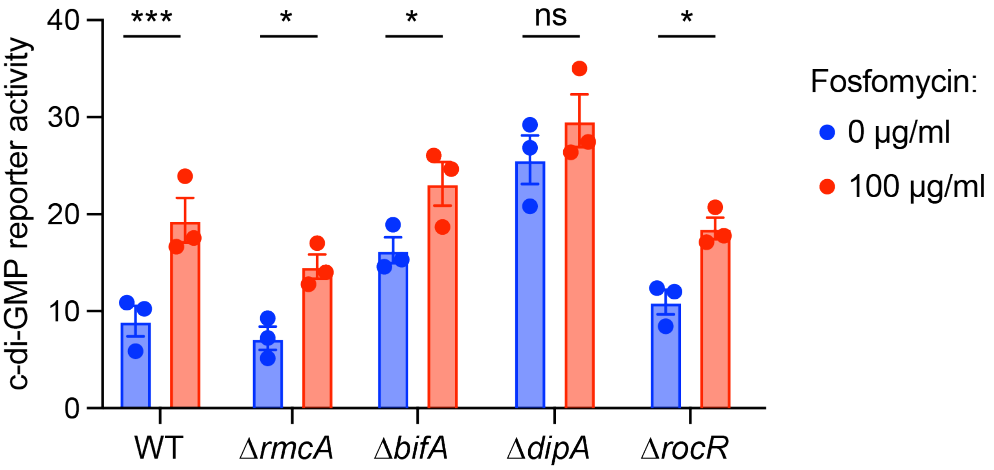


**Supplementary Fig. 8. Effects of fosfomycin on selective c-di-GMP PDE mutants.** c-di-GMP reporter activity in response to 100 µg/ml fosfomycin in indicated strains. **p* < 0.05, ***p* < 0.01, ****p* < 0.001, determined by two-way ANOVA with Sidak's multiple comparisons test.

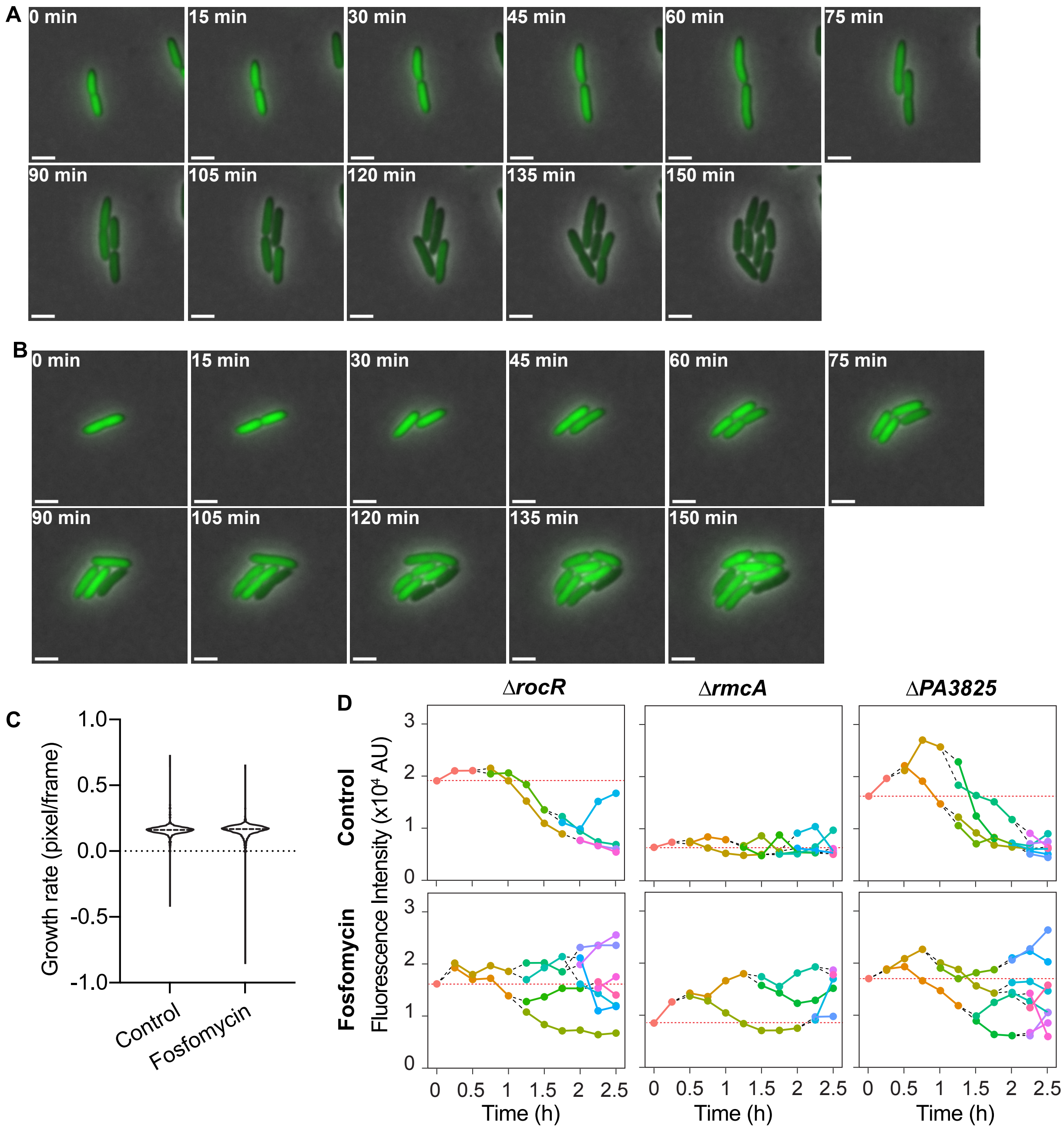


**Supplementary Fig. 9. cdGreen fluorescence and growth on the agarose pad.** (A) Micrographs of cells grown on the control agarose pad, corresponding to the lineage analysis shown in Fig. 4C. (B) Micrographs of cells grown on the agarose pad containing 100 µg/ml fosfomycin, corresponding to the lineage analysis shown in Fig. 4D. (C) Violin plot showing bacterial growth rates on agarose pads with or without fosfomycin. No significant difference was observed, determined by a two-tailed t test. (D) Representative cell lineages of indicated strains in control and fosfomycin treated conditions. Points with connecting lines indicate measurement of the same cell over time. Black dash lines mark cell division events, and orange dotted lines represent cdGreen2 fluorescence at the inoculum.

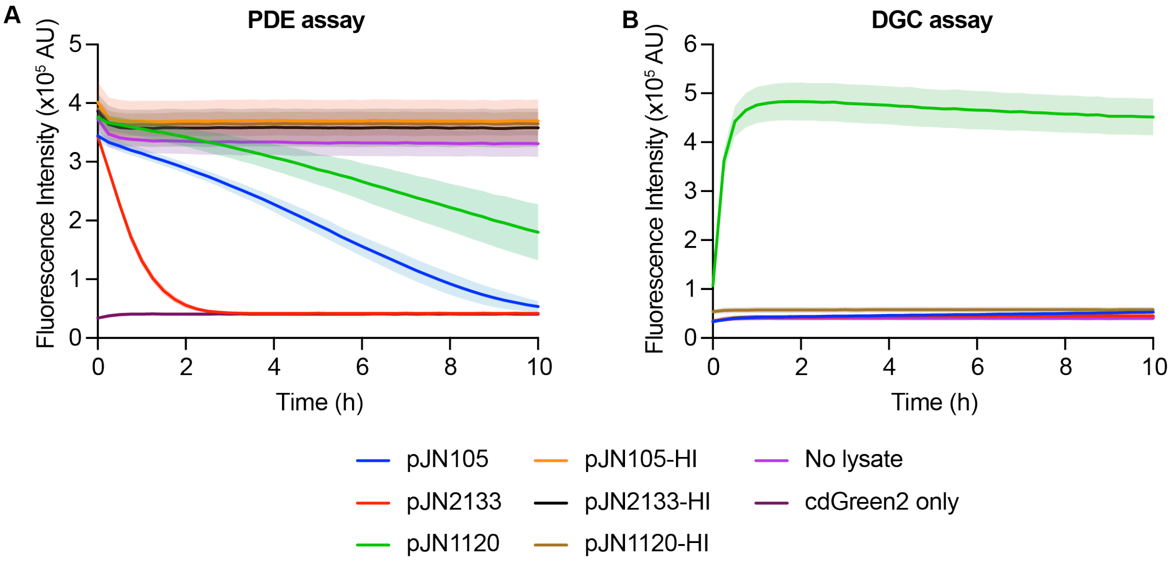


**Supplementary Fig. 10. Validation of *in vitro* c-di-GMP PDE and DGC assay.** In vitro PDE or DGC assays were performed using purified cdGreen2 and bacterial cell lysates containing either an empty vector plasmid (pJN105), a plasmid overexpressing the PDE PA2133 (pJN2133), or a plasmid overexpressing the DGC PA1120 (pJN1120). Heat inactivated (HI) lysates, no-lysate, and no-substrate/no-lysate were included as controls. Data show mean ± SEM of 3 independent experiments, demonstrating PA2133-dependent PDE activity (A) and PA1120-dependent DGC activity (B).

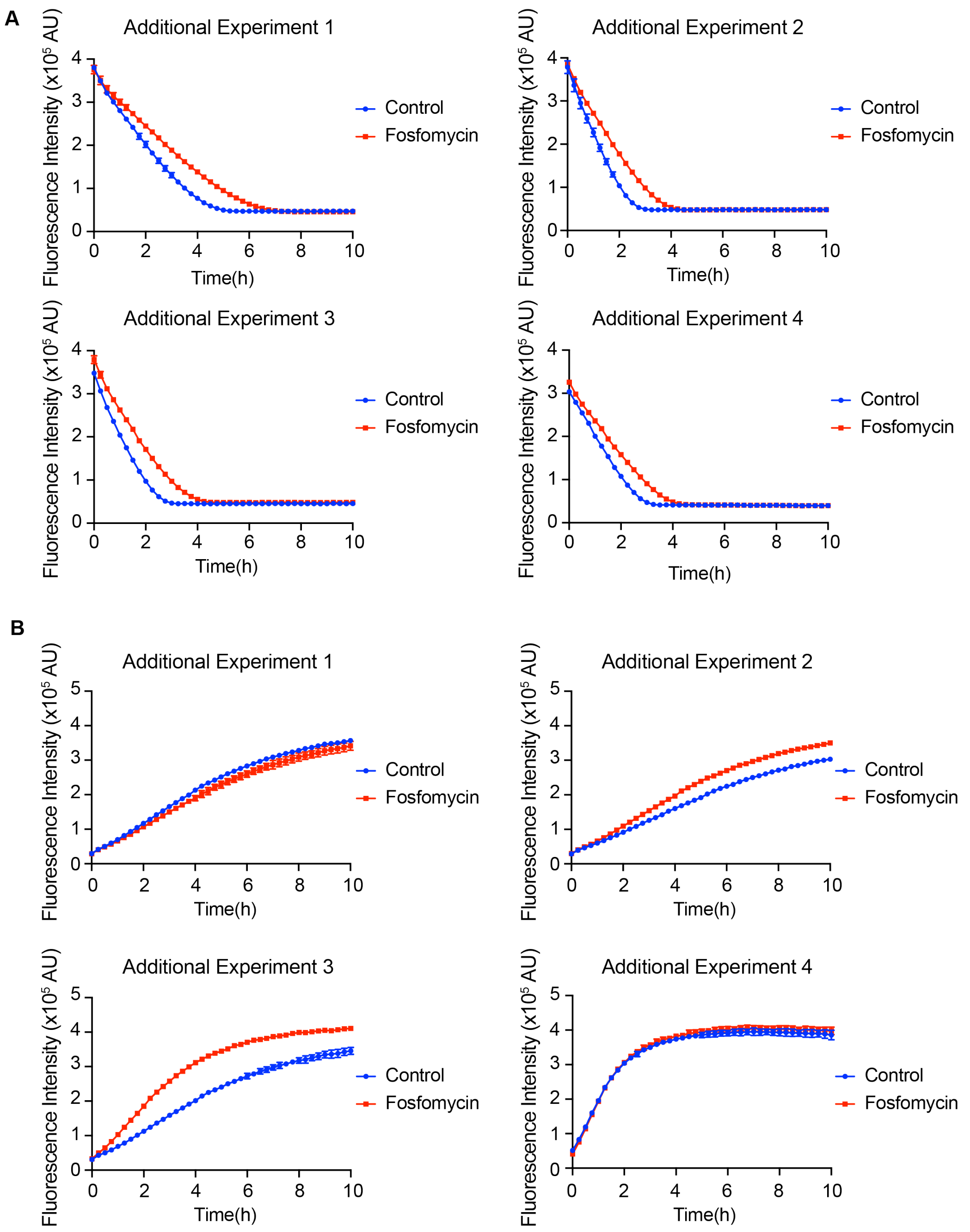


**Supplementary Fig. 11. Additional biological replicates for the PDE and DGC activity assays.** Comparison of WT and fosfomycin-treated lysates show significant variations in the kinetics of c-di-GMP degradation and synthesis, although fosfomycin-grown lysate consistently exhibits reduced PDE activity. This figure shows the additional experiments complementing Fig. 6, evaluating PDE (A) and DGC (B) activity. Each graph is an independent experiment.

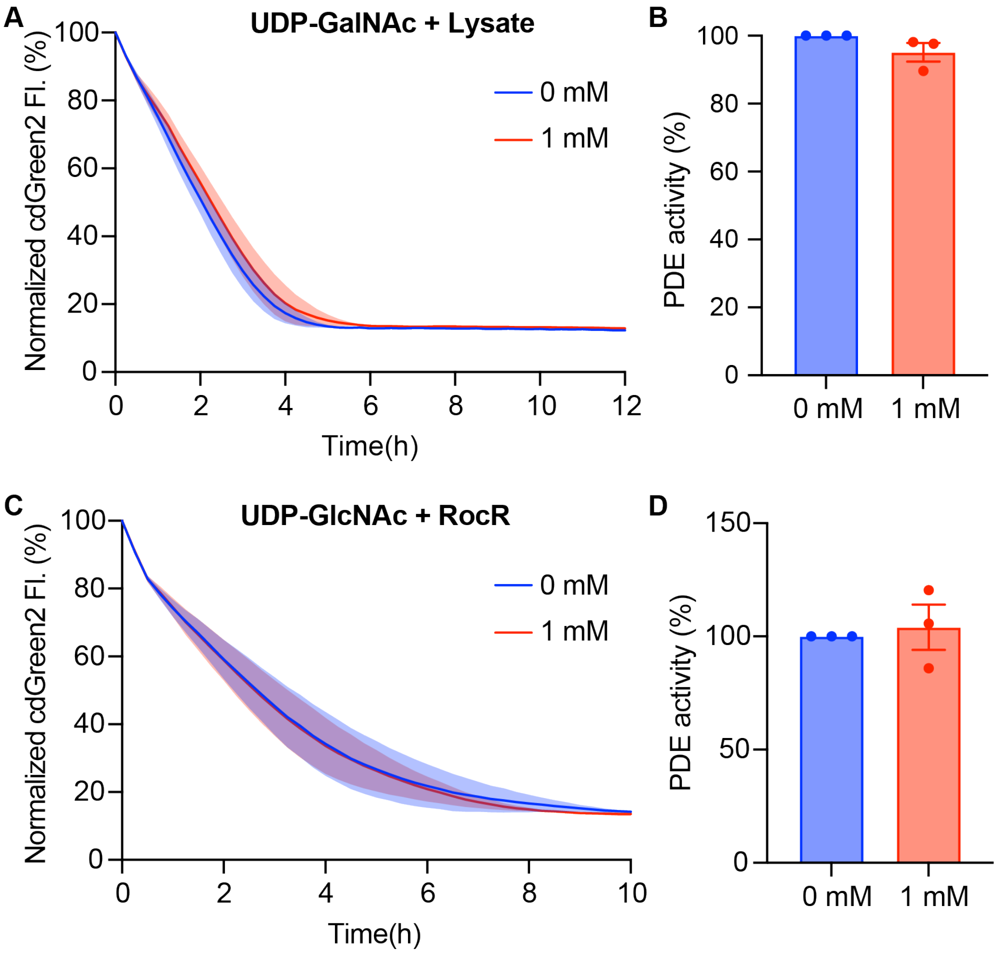


**Supplementary Fig. 12. UDP-GalNAc minimally affect c-di-GMP PDE activity *in vitro*.** (A) Kinetics of c-di-GMP degradation with cell lysates treated with or without 1 mM UDP-GalNAc. Data shown mean ± SEM of three independent experiments. (B) Quantification of c-di-GMP PDE activity derived from (A). Data are normalized to the untreated condition in each experiment and represent show mean ± SEM of three independent experiments. (C) Kinetics of c-di-GMP degradation with purified RocR treated with or without 1 mM UDP-GlcNAc. Data shown mean ± SEM of three independent experiments. (D) Quantification of c-di-GMP PDE activity derived from (C). Data are normalized to the untreated condition in each experiment and represent show mean ± SEM of three independent experiments.

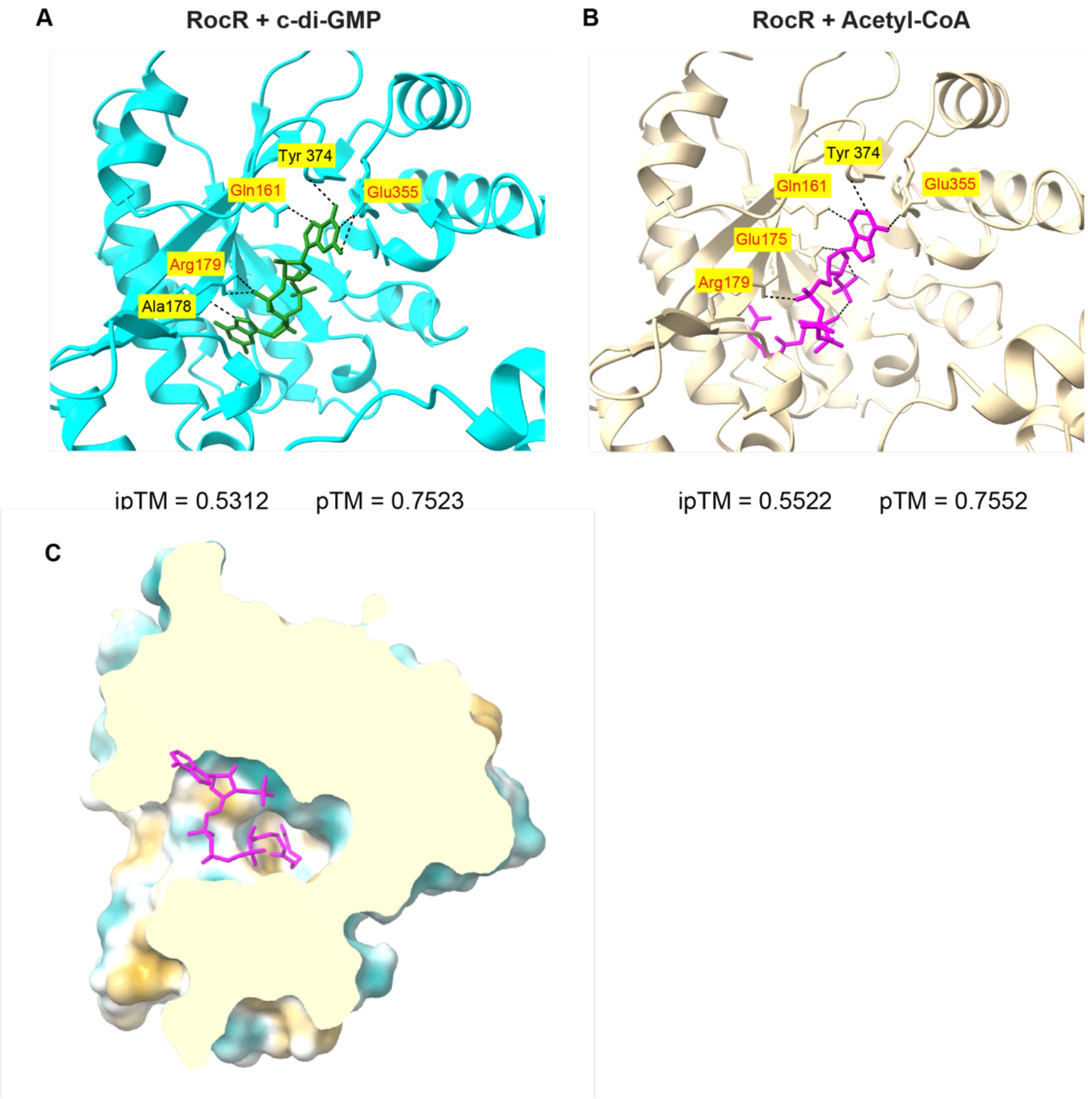


**Supplementary Fig. 13. AlphaFold3 prediction of c-di-GMP or acetyl-CoA interaction with RocR.** (A-B) Predicted hydrogen bonds formed between c-di-GMP (A) or acetyl-CoA (B) and RocR. Red indicates conserved residues among EAL domains across species. (C) Predicted acetyl-CoA docking into RocR, highlighting the acyl chain into a groove with mid-high hydrophobicity. The coloring shows relative surface hydrophobicity of RocR.

**Supplementary Table 1. List of molecules screened for c-di-GMP reporter activity.**

| Name | Source | Function | Class | Reference |
|---|---|---|---|---|
| Sodium dodecyl sulfate (SDS) | Fisher BioReagents BP166 | Anionic detergent; solubilizes membrane | General | (1) |
| Ethanol | Decon Labs 04355223 | Influences membrane properties | General | (2) |
| Triton X-100 | Sigma-Aldrich T8787 | Nonionic detergent; solubilizes membrane | General | (1) |
| Glycerol | Sigma-Aldrich G7893 | Osmolyte; influences osmotic pressure | General | |
| Urea | Fisher Scientific A12360 | Denatures proteins | General | |
| $H_2O_2$ | Supelco HX0635 | Induces oxidative stress | General | |
| Nigericin | Sigma-Aldrich N7143 | $H^+/K^+$ antiporter; Dissipates proton gradient | Membrane | (3) |
| Oligomycin | Sigma-Aldrich 495455 | ATP synthase inhibitor; Increases membrane potential | Membrane | (3) |
| Valinomycin | Sigma-Aldrich V0627 | $K^+$ ionophore; Reduces membrane potential | Membrane | (3) |
| NaCl | Fisher BioReagents BP358 | Influences osmotic pressure | Membrane | |
| Benzyl Alcohol | Sigma-Aldrich 305197 | Increases membrane fluidity | Membrane | (4) |
| Fosfomycin | Sigma-Aldrich P5396 | Inhibits murA | Cell Wall | (5) |
| D-cycloserine | Sigma-Aldrich 30020 | D-alanine analog; blocks D-ala incorporation | Cell Wall | (6) |
| Tunicamycin | Sigma-Aldrich T7765 | Inhibits MraY | Cell Wall | (7) |

| | | | | |
|---|---|---|---|---|
| Ramoplanin | Cayman Chemical 28189 | Sequesters lipid I and lipid II | Cell Wall | (8) |
| Moenomycin | Cayman Chemical 15506 | Inhibits glycosyltransferase | Cell Wall | (9) |
| Carbenicillin | Sigma-Aldrich C1389 | Penicillin group of β-lactam; inhibits penicillin binding proteins (PBPs) | Cell Wall | (10) |
| Ceftazidime | Fisher Scientific J6646003 | Cephalosporin group of β-lactam; inhibits PBPs | Cell Wall | (10) |
| Meropenem | Supelco PHR1772 | Carbapenem group of β-lactam; inhibits PBPs | Cell Wall | (10) |
| Colistin | Research Products International C70700 | Bind LPS; disrupts outer membrane | LPS | (11) |
| Polymyxin B | Research Products International P40160 | Bind LPS; disrupts outer membrane | LPS | (11) |
| Bacitracin | Sigma-Aldrich B5150 | Inhibits dephosphorylation of lipid carrier undecaprenyl pyrophosphate | Fatty Acid | (12) |
| Fosmidomycin | Cayman Chemical 16000 | Inhibits isoprenoid biosynthesis | Fatty Acid | (13) |
| Triclosan | Sigma-Aldrich 72779 | Inhibits fatty acid biosynthesis | Fatty Acid | (14) |
| A22 | Sigma-Aldrich SML0471 | Inhibits MreB | Cytoskeleton | (15) |
| Trimethoprim | Sigma-Aldrich T7883 | Inhibits thymidine synthesis | DNA | (16) |
| Ciprofloxacin | Sigma-Aldrich 17850 | Inhibits DNA gyrase and topoisomerase IV | DNA | (17) |

| | | | | |
|---|---|---|---|---|
| Kanamycin | TOKU-E K008 | Binds to 30s rRNA and inhibit translation | Protein | (18) |
| Tetracycline | Fisher BioReagents BP912 | Inhibits tRNA binding to ribosome | Protein | (19) |

**Supplementary Table 2. Bacterial strains used in this study.**

| No. | Strain | Description | Source |
|---|---|---|---|
| *E. coli* | | | |
| | DH5a | Strain for cloning. | NEB |
| | BL21(DE3) | Strain for protein purification | NEB |
| *P. aeruginosa* | | | |
| XZ1 | PAO1 | PAO1 (Parsek) wild-type. | (20) |
| XZ68 | Δ*wspR*Δ*sadC*Δ*siaD* | PAO1 with markerless deletions of *wspR, sadC,* and *siaD.* | (21) |
| XZ658 | Δ*pel*Δ*psl*Δ*cdrA* | PAO1 with markerless deletions of *pelA*, *pslBCD*, and *cdrA*. | (22) |
| XZ659 | Tn7::*araBAD-dCas9* | PAO1 with arabinose-inducible deactivated Cas9 in the mini-Tn7 site. | O'Malley et al., 2025 |
| XZ821 | PAO1L | PAO1 (Lausanne) wild-type. | (23) |
| XZ822 | ΔPA0338 | PAO1L mutant of ΔPA0338. | (23) |
| XZ823 | ΔPA0847 | PAO1L mutant of ΔPA0847. | (23) |
| XZ824 | ΔPA3343 | PAO1L mutant of ΔPA3343 (hsbD). | (23) |
| XZ825 | ΔPA3702 | PAO1L mutant of ΔPA3702 (wspR). | (23) |
| XZ826 | ΔPA4332 | PAO1L mutant of ΔPA4332 (sadC). | (23) |
| XZ827 | ΔPA2072 | PAO1L mutant of ΔPA2072. | (23) |
| XZ828 | ΔPA4367 | PAO1L mutant of ΔPA4367 (bifA). | (23) |
| XZ829 | ΔPA0290 | PAO1L mutant of ΔPA0290. | (23) |
| XZ830 | ΔPA3177 | PAO1L mutant of ΔPA3177. | (23) |
| XZ831 | ΔPA4396 | PAO1L mutant of ΔPA4396. | (23) |
| XZ832 | ΔPA4843 | PAO1L mutant of ΔPA4843 (gcbA). | (23) |
| XZ833 | ΔPA1851 | PAO1L mutant of ΔPA1851. | (23) |
| XZ834 | ΔPA2818 | PAO1L mutant of ΔPA2818 (arr). | (23) |
| XZ835 | ΔPA3825 | PAO1L mutant of ΔPA3825. | (23) |
| XZ836 | ΔPA2567 | PAO1L mutant of ΔPA2567. | (23) |
| XZ837 | ΔPA5487 | PAO1L mutant of ΔPA5487 (dgcH). | (23) |
| XZ838 | ΔPA5295 | PAO1L mutant of ΔPA5295 (proE). | (23) |
| XZ839 | ΔPA3258 | PAO1L mutant of ΔPA3258. | (23) |
| XZ840 | ΔPA1727 | PAO1L mutant of ΔPA1727 (mucR). | (23) |
| XZ841 | ΔPA2771 | PAO1L mutant of ΔPA2771. | (23) |

| | | | |
|---|---|---|---|
| XZ842 | ΔPA1107 | PAO1L mutant of ΔPA1107 (roeA). | (23) |
| XZ843 | ΔPA1120 | PAO1L mutant of ΔPA1120 (tpbB). | (23) |
| XZ844 | ΔPA2870 | PAO1L mutant of ΔPA2870. | (23) |
| XZ845 | ΔPA4929 | PAO1L mutant of ΔPA4929. | (23) |
| XZ846 | ΔPA4959 | PAO1L mutant of ΔPA4959 (fimX). | (23) |
| XZ847 | ΔPA0861 | PAO1L mutant of ΔPA0861 (rbdA). | (23) |
| XZ848 | ΔPA1181 | PAO1L mutant of ΔPA1181. | (23) |
| XZ849 | ΔPA1433 | PAO1L mutant of ΔPA1433. | (23) |
| XZ850 | ΔPA3311 | PAO1L mutant of ΔPA3311 (nbdA). | (23) |
| XZ851 | ΔPA4601 | PAO1L mutant of ΔPA4601 (morA). | (23) |
| XZ852 | ΔPA5017 | PAO1L mutant of ΔPA5017 (dipA). | (23) |
| XZ853 | ΔPA2200 | PAO1L mutant of ΔPA2200. | (23) |
| XZ854 | ΔPA3947 | PAO1L mutant of ΔPA3947 (rocR). | (23) |
| XZ855 | ΔPA2572 | PAO1L mutant of ΔPA2572. | (23) |
| XZ856 | ΔPA4781 | PAO1L mutant of ΔPA4781. | (23) |
| XZ857 | ΔPA0285 | PAO1L mutant of ΔPA0285 (pipA). | (23) |
| XZ858 | ΔPA2133 | PAO1L mutant of ΔPA2133. | (23) |
| XZ859 | ΔPA5442 | PAO1L mutant of ΔPA5442. | (23) |
| XZ860 | ΔPA0169 | PAO1L mutant of ΔPA0169 (siaD). | (23) |
| XZ861 | ΔPA0575 | PAO1L mutant of ΔPA0575 (rmcA). | (23) |
| XZ862 | ΔPA4108 | PAO1L mutant of ΔPA4108. | (23) |

**Supplementary Table 3. Plasmids used in this study.**

| No. | Name | Description | Source |
|---|---|---|---|
| XZ23 | pBBR1MCS-5 | Cloning vector. GmR. | (24) |
| XZ44 | pEX18Gm | Allelic exchange vector. GmR. | (25) |
| XZ198 | pBBR1MCS5-Ppaqa-mScarlet(ASV)-tet2-PcdrA-mGL(ASV)-t0t1-PrpoD-SCFP3A(ASV) | Tri-color reporter used a template to generate c-di-GMP reporter plasmid. | (21) |
| XZ197 | pBBR1MCS5-PcdrA-mGL(ASV)-PrpoD-mScarlet-I(ASV) | c-di-GMP reporter plasmid with a constitutive promoter control. GmR. | This study |
| XZ654 | pBBR1MCS5-PcdrA-mGL(ASV)-tet2-PrpoD-mScarlet-I(ASV)-gEV | c-di-GMP reporter plasmid with the constitutive promoter but no sgRNA. GmR. | This study |
| XZ665 | pBBR1MCS5-PcdrA-mGL(ASV)-tet2-PrpoD-mScarlet-I(ASV)-gmurA | c-di-GMP reporter plasmid with a constitutive promoter driving the expression of sgRNA against murA. GmR. | This study |
| XZ667 | pBBR1MCS5-PcdrA-mGL(ASV)-tet2-PrpoD-mScarlet-I(ASV)-gmurF | c-di-GMP reporter plasmid with a constitutive promoter driving the expression of sgRNA against murF. GmR. | This study |
| XZ668 | pBBR1MCS5-PcdrA-mGL(ASV)-tet2-PrpoD-mScarlet-I(ASV)-gmurJ | c-di-GMP reporter plasmid with a constitutive promoter driving the expression of sgRNA against murJ. GmR. | This study |
| XZ671 | pBBR1MCS5-PcdrA-mGL(ASV)-tet2-PrpoD-mScarlet-I(ASV)-guppS | c-di-GMP reporter plasmid with a constitutive promoter driving the expression of sgRNA against uppS. GmR. | This study |
| AS93 | pBBR1MCS5-PlacP1-mGL(ASV)-tet2-PrpoD-mScarlet-I(ASV)-gmurB | c-di-GMP reporter plasmid with a constitutive promoter driving the expression of sgRNA against murB. GmR. | This study |
| AS131 | pBBR1MCS5-PlacP1-mGL(ASV)-tet2-PrpoD-mScarlet-I(ASV)-glpxA | c-di-GMP reporter plasmid with a constitutive promoter driving the expression of sgRNA against lpxA. GmR. | This study |
| AS134 | pBBR1MCS5-PlacP1-mGL(ASV)-tet2-PrpoD-mScarlet-I(ASV)-gglmU | c-di-GMP reporter plasmid with a constitutive promoter driving the expression of sgRNA against glmU. GmR. | This study |

| | | | |
|---|---|---|---|
| XZ807 | pBBR15.2-2H12.D11opt | pBBR1 plasmid containing the cdGreen reporter. GmR. | (26) |
| XZ809 | pET28-2H12.D11 | pET28a plasmid for cdGreen purification. KanR. | (26) |
| XZ66 | pJN105 | pBBR1 containing the *araC*-P*BAD* cassette. GmR. | (27) |
| XZ881 | pJN2133 | pJN105 with arabinose-inducible PA2133. | (28) |
| XZ882 | pJN1120 | pJN105 with arabinose-inducible PA1120. | (29) |

**Supplementary Table 4. Primers used in this study.**

| No. | Sequence | Product |
|---|---|---|
| oXZ213 | AAAAGACAATGAACCCCCGCTCGAGcattgtcggttttttgacgg | PcdrA-mGL(ASV) |
| oXZ206 | CCGTTTCCACGGTGTGCGTCCATGGGAATTCttaaactgatgcagcgtagttttc | PcdrA-mGL(ASV) |
| oXZ225 | ctcgagcgggggttcattg | T2Te |
| oXZ229 | GCATGCGCGGATTTGAAC | T2Te |
| oXZ216 | CGCAACGTTCAAATCCGCGCATGCatacctacgcccagttgctg | PrpoD |
| oXZ21 | aacaccctatccactgaaggtc | PrpoD |
| oXZ22 | gaccttcagtggatagggtgttatgagtaaaggagaagctgtgattaaag | mScarlet-I(ASV) |
| oXZ230 | ATAGGGCGAATTGGAGCTCCCCGGGttaaactgatgcagcgtagttttc | mScarlet-I(ASV) |
| MO172 | ACGCTGCATCAGTTTAACCCTTGACAGCTAGCTCAGTCCT | PspeI-sgRNA |
| MO173 | AGGGCGAATTGGAGCTCCCCCAACTGTTGGGAAGGGCGAT | PspeI-sgRNA |
| oXZ434 | ACTAGTATTATACCTAGGACTGAGCTAGC | gRNA |
| oXZ436 | GCTCAGTCCTAGGTATAATACTAGTgccggaaatgcggatttcgccatcgGTTTTAGAGCTAGAAATAGCAAGTTAAAATAAGGC | gRNA (murA) |
| oXZ438 | GCTCAGTCCTAGGTATAATACTAGTaagacggcgtcctcgccgatcaggcGTTTTAGAGCTAGAAATAGCAAGTTAAAATAAGGC | gRNA (murF) |
| oXZ439 | GCTCAGTCCTAGGTATAATACTAGTccgaggacccgcgacagcatggtgaGTTTTAGAGCTAGAAATAGCAAGTTAAAATAAGGC | gRNA (murJ) |
| oXZ442 | GCTCAGTCCTAGGTATAATACTAGTcgtccatgataatggccacgtggcgGTTTTAGAGCTAGAAATAGCAAGTTAAAATAAGGC | gRNA (uppS) |
| oXZ503 | GCTCAGTCCTAGGTATAATACTAGTgcacgtcgatgccgaaggtgttataGTTTTAGAGCTAGAAATAGCAAGTTAAAATAAGGC | gRNA (murB) |
| oXZ512 | GCTCAGTCCTAGGTATAATACTAGTtggcgctcggatcgatgatggcgcgGTTTTAGAGCTAGAAATAGCAAGTTAAAATAAGGC | gRNA (lpxA) |
| oXZ582 | agtcgtattacgcgcgc | double CRISPRi plasmid backbone |
| oXZ581 | cactatagggcgaattggagc | double CRISPRi plasmid backbone |
| oXZ579 | GCTCCAATTCGCCCTATAGTGcccttgacagctagctcag | second gRNA |
| oXZ580 | TGAGCGCGCGTAATACGACTgagctcccccaactgttg | second gRNA |

### REFERENCES


1. A. Helenius, K. Simons, Solubilization of membranes by detergents. *Biochimica et Biophysica Acta (BBA) - Reviews on Biomembranes* **415**, 29–79 (1975).
2. L. O. Ingram, T. M. Buttke, "Effects of Alcohols on Micro-Organisms" in (1985), pp. 253–300.
3. F. M. Harold, "Antimicrobial Agents and Membrane Function" in (1969), pp. 45–104.
4. A. Chabanel, R. E. Abbott, S. Chien, D. Schachter, Effects of benzyl alcohol on erythrocyte shape, membrane hemileaflet fluidity and membrane viscoelasticity. *Biochimica et Biophysica Acta (BBA) - Biomembranes* **816**, 142–152 (1985).
5. L. L. Silver, Fosfomycin: Mechanism and Resistance. *Cold Spring Harb Perspect Med* **7**, a025262 (2017).
6. F. C. Neuhaus, J. L. Lynch, The Enzymatic Synthesis of D-Alanyl-D-alanine. III. On the Inhibition of D-Alanyl-D-alanine Synthetase by the Antibiotic D-Cycloserine [*]. *Biochemistry* **3**, 471–480 (1964).
7. J. K. Hakulinen, *et al.*, MraY-antibiotic complex reveals details of tunicamycin mode of action. *Nat Chem Biol* **13**, 265–267 (2017).
8. P. Cudic, *et al.*, Complexation of peptidoglycan intermediates by the lipoglycodepsipeptide antibiotic ramoplanin: Minimal structural requirements for intermolecular complexation and fibril formation. *Proceedings of the National Academy of Sciences* **99**, 7384–7389 (2002).
9. T.-J. R. Cheng, *et al.*, Domain requirement of moenomycin binding to bifunctional transglycosylases and development of high-throughput discovery of antibiotics. *Proceedings of the National Academy of Sciences* **105**, 431–436 (2008).
10. K. Bush, P. A. Bradford, β-Lactams and β-Lactamase Inhibitors: An Overview. *Cold Spring Harb Perspect Med* **6**, a025247 (2016).
11. R. E. Hancock, Peptide antibiotics. *The Lancet* **349**, 418–422 (1997).
12. G. Siewert, J. L. Strominger, BACITRACIN: AN INHIBITOR OF THE DEPHOSPHORYLATION OF LIPID PYROPHOSPHATE, AN INTERMEDIATE IN THE BIOSYNTHESIS OF THE PEPTIDOGLYCAN OF BACTERIAL CELL WALLS. *Proceedings of the National Academy of Sciences* **57**, 767–773 (1967).
13. E. S. McKenney, *et al.*, Lipophilic Prodrugs of FR900098 Are Antimicrobial against Francisella novicida In Vivo and In Vitro and Show GlpT Independent Efficacy. *PLoS One* **7** (2012).
14. R. J. Heath, *et al.*, Mechanism of Triclosan Inhibition of Bacterial Fatty Acid Synthesis. *Journal of Biological Chemistry* **274**, 11110–11114 (1999).
15. Z. Gitai, N. A. Dye, A. Reisenauer, M. Wachi, L. Shapiro, MreB actin-mediated segregation of a specific region of a bacterial chromosome. *Cell* **120**, 329–341 (2005).

16. P. A. Masters, T. A. O'Bryan, J. Zurlo, D. Q. Miller, N. Joshi, Trimethoprim-Sulfamethoxazole Revisited. *Arch Intern Med* **163**, 402 (2003).

17. K. J. Aldred, R. J. Kerns, N. Osheroff, Mechanism of Quinolone Action and Resistance. *Biochemistry* **53**, 1565–1574 (2014).

18. B. François, *et al.*, Crystal structures of complexes between aminoglycosides and decoding A site oligonucleotides: Role of the number of rings and positive charges in the specific binding leading to miscoding. *Nucleic Acids Res* **33**, 5677–5690 (2005).

19. C. U. Chukwudi, rRNA binding sites and the molecular mechanism of action of the tetracyclines. *Antimicrob Agents Chemother* [Preprint] (2016).

20. B. R. Borlee, *et al.*, Pseudomonas aeruginosa uses a cyclic-di-GMP-regulated adhesin to reinforce the biofilm extracellular matrix. *Mol Microbiol* **75**, 827–842 (2010).

21. X. Zheng, *et al.*, The surface interface and swimming motility influence surface-sensing responses in Pseudomonas aeruginosa. *Proc Natl Acad Sci U S A* **121**, e2411981121 (2024).

22. C. Reichhardt, C. Wong, D. Passos da Silva, D. J. Wozniak, M. R. Parsek, CdrA Interactions within the Pseudomonas aeruginosa Biofilm Matrix Safeguard It from Proteolysis and Promote Cellular Packing. *mBio* **9** (2018).

23. K. Eilers, *et al.*, Phenotypic and integrated analysis of a comprehensive Pseudomonas aeruginosa PAO1 library of mutants lacking cyclic-di-GMP-related genes. *Front Microbiol* **13**, 949597 (2022).

24. M. E. Kovach, R. W. Phillips, P. H. Elzer, R. M. Roop, K. M. Peterson, pBBR1MCS: a broad-host-range cloning vector. *Biotechniques* **16**, 800–2 (1994).

25. L. R. Hmelo, *et al.*, Precision-engineering the Pseudomonas aeruginosa genome with two-step allelic exchange. *Nat Protoc* **10**, 1820–1841 (2015).

26. A. Kaczmarczyk, *et al.*, A genetically encoded biosensor to monitor dynamic changes of c-di-GMP with high temporal resolution. *Nat Commun* **15**, 3920 (2024).

27. J. R. Newman, C. Fuqua, Broad-host-range expression vectors that carry the l-arabinose-inducible Escherichia coli araBAD promoter and the araC regulator. *Gene* **227**, 197–203 (1999).

28. J. W. Hickman, D. F. Tifrea, C. S. Harwood, A chemosensory system that regulates biofilm formation through modulation of cyclic diguanylate levels. *Proceedings of the National Academy of Sciences* **102**, 14422–14427 (2005).

29. J. W. Hickman, C. S. Harwood, Identification of FleQ from Pseudomonas aeruginosa as a c-di-GMP-responsive transcription factor. *Mol Microbiol* **69**, 376–389 (2008).
